# Protein Language Models Trained on Biophysical Dynamics Inform Mutation Effects

**DOI:** 10.1101/2024.10.11.617911

**Authors:** Chao Hou, Haiqing Zhao, Yufeng Shen

## Abstract

Structural dynamics are fundamental to protein functions and mutation effects. Current protein deep learning models are predominantly trained on sequence and/or static structure data, which often fail to capture the dynamic nature of proteins. To address this, we introduce SeqDance and ESMDance, two protein language models trained on dynamic biophysical properties derived from molecular dynamics simulations and normal mode analyses of over 64,000 proteins. Both models can be directly applied to predict dynamic properties of unseen ordered and disordered proteins. SeqDance, trained from scratch, has attentions that capture dynamic interaction and co-movement between residues, and its embeddings encode rich representations of protein dynamics that can be further utilized to predict conformational properties beyond the training tasks via transfer learning. SeqDance predicted dynamic property changes reflect mutation effect on protein folding stability. ESMDance, built upon ESM2 (Evolutionary Scale Model II) outputs, substantially outperforms ESM2 in zero-shot prediction of mutation effects for designed and viral proteins which lack evolutionary information. Together, SeqDance and ESMDance offer a new framework for integrating protein dynamics into language models, enabling more generalizable predictions of protein behavior and mutation effects.

**Significance Statement:** The sequence—structure (ensemble)—function relationship is central to biology. Protein dynamics in the structure ensemble play a decisive role in determining function and mutation effects, and are widely used to study thermodynamics, folding pathways, and dynamic interactions of ordered proteins, as well as the conformational variability of intrinsically disordered proteins. However, current state-of-the-art protein deep learning models, such as AlphaFold2,3 and ESM, focus on static structures and sequences, which failed to directly capture protein dynamics. Here, we address this gap by developing protein language models to learn dynamic properties of over 64,000 proteins. We show that the model’s Transformer attentions capture protein dynamic interactions, and our model can be applied to predict conformational properties and mutation effects.

## Introduction

Protein deep learning models have made significant progresses in predicting 3D structure (e.g., AlphaFold(1, 2), RoseTTAfold(3), and ESMFold(4)), protein function(5), stability(6), localization(7), and interactions(8). A central challenge in developing effective models lies in the selection of information sources that models can interpret and learn from. Broadly, these information can be classified into two main types: evolutionary and biophysical(9).

Evolutionary information is derived from protein homologous sequences across species that have been shaped by similar selection pressures. Multiple sequence alignments (MSAs) of homologs contain information of conserved motifs and co-evolutionary pairs. Recently, protein language models (pLMs) like ESM1,2(4, 10), ProtTrans(11), and ProGen(12), trained on large-scale protein sequence datasets, have emerged as powerful tools to implicitly learn evolutionary information(13, 14). pLMs learn rich representations of evolutionary information and perform well across diverse tasks, making them common baselines for evaluating new models. MSAs and pLMs have been successfully applied to predict 3D structures(1, 4), predict pathogenic mutations(15, 16), identify signal peptides(6), and even generate novel proteins(12). However, the effectiveness of evolutionary information relies on the quality and quantity of available homologous sequences. For instance, the confidences of structure predictions correlate to the number of homologous sequences(1, 4). As a result, evolutionary information is less effective for intrinsically disordered regions (IDRs), designed proteins, rapidly evolving viral proteins, immune proteins, and those from under-studied species like extremophiles(17), where homologous sequences are either sparse or highly divergent. Moreover, evolutionary patterns in homologs are the outcomes of protein functions rather than the underlying causes (Figure 1A). Overreliance on evolutionary information may bias models toward correlational patterns rather than learning fundamental principles governing protein behaviors.

**Figure 1:**
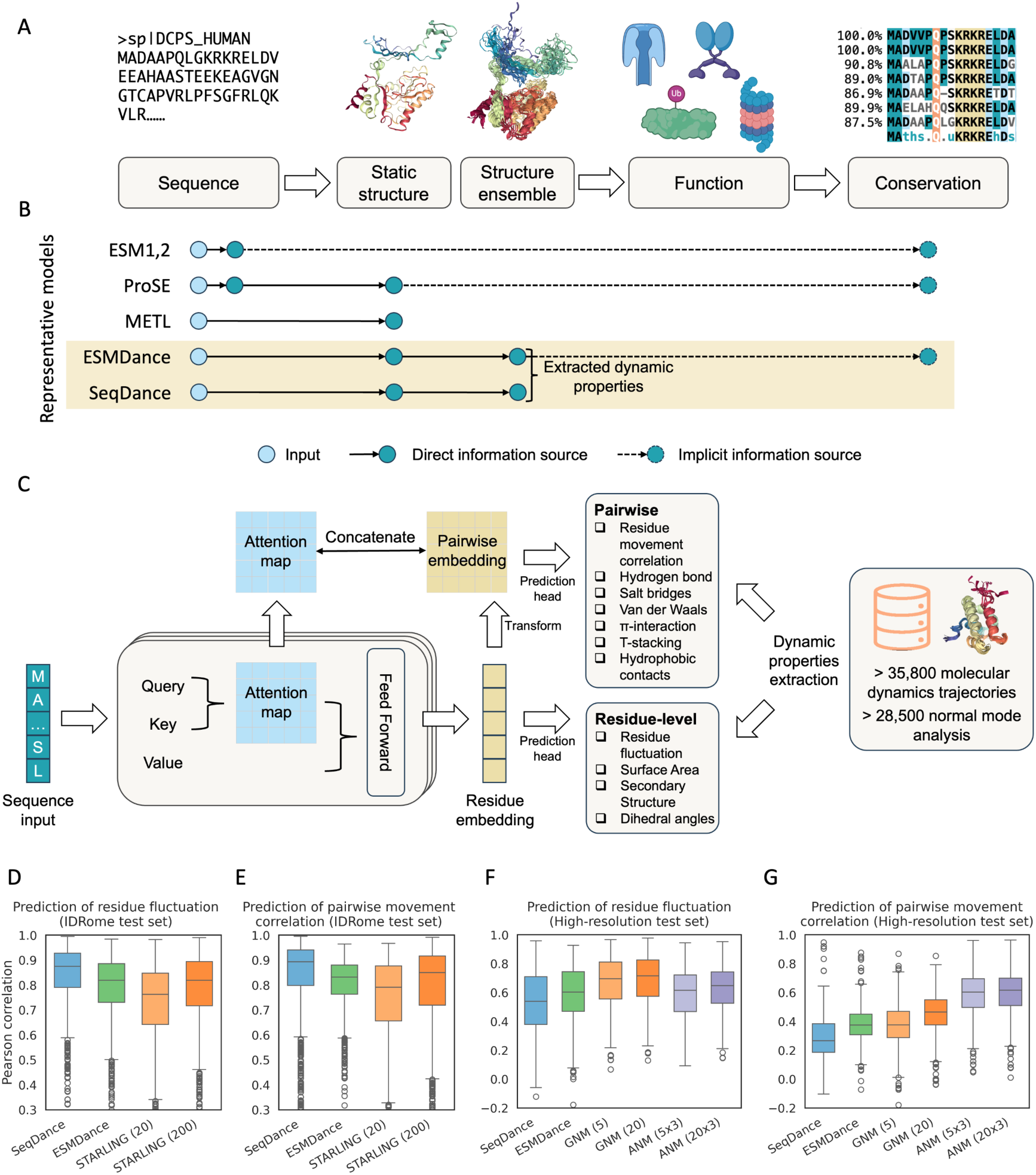
Information flow in protein study, representative protein language models, model pre-training and performance on test set**. A.** Illustration of the “sequence - structure ensemble - function - evolution” paradigm. The sequence is the basis of a protein, which folds into a structural ensemble to perform its function. Functionally important region exhibits conserved patterns across homologs. **B**. Representative protein language models (pLMs) and their information sources. ESM1 and 2 were trained to predict masked residues and implicitly learned evolutionary information. ProSE was trained to predict masked residues, pairwise contact in static structures, and structure similarity. METL was trained to predict biophysical terms calculated from static structures. SeqDance and ESMDance were trained on protein dynamic properties from molecular dynamics (MD) simulations, experimental data, and normal mode analysis (NMA) of static structures. **C**. Diagram of the pre-training process. SeqDance and ESMDance take a protein sequence as input and predict residue-level and pairwise dynamic properties, which are extracted from structures ensembles and NMA. Both models use a Transformer encoder architecture identical to ESM2-35M, consisting of 12 layers with 20 heads per layer and an embedding dimension of 480. Linear layers are applied to the residue embeddings to predict residue-level properties. For pairwise property prediction, pairwise embeddings, constructed from residue embeddings, are concatenated with attention maps, a linear layer is then applied to predict pairwise properties. **D-E**. Comparison of SeqDance, ESMDance, and STARLING on 1,249 test IDRs (filtered to sequences ≤384 residues, since STARLING cannot be applied to longer IDRs). STARLING was used to generate 20 or 200 conformations for each IDR. **F-G**. Comparison of SeqDance, ESMDance, and two NMA methods on 404 test high-resolution ordered proteins (filtered to sequences ≤1024 residues, since SeqDance and ESMDance were trained with max length of 1024). The low-frequency 5 or 20 modes from Gaussian Network Model (GNM) and 5×3 or 20×3 modes from Anisotropic Network Model (ANM) were used for each protein (see Methods for details). In those boxplots, the box extends from the first quartile to the third quartile of the data, with a line at the median. The whiskers extend from the box to the farthest data point lying within 1.5 times the inter-quartile range from the box.

Biophysical information, primarily derived from protein structures, circumvents the limitations of evolutionary information. Predicting protein behaviors from biophysical information aligns with the central paradigm that sequence determines structure, and structure governs function (Figure 1A), thereby enabling models to generalize more broadly across the protein universe. Biophysical information has been applied to a wide range of biological questions. For example, predicted structures have been used to predict mutation effects and functional sites(18). MaSIF(8) utilized biophysical and geometric features on protein surface to study protein interactions. PScore(19) was trained to predict Pi-interactions derived from PDB(20) structures and can be used to identify phase-separating proteins(21). Additionally, there have been initiatives to integrate biophysical information into pLMs. ProSE(9) was trained to predict masked residues, contacts within static structures, and structural similarities, while METL(22) was developed to predict biophysical properties derived from static structures (Figure 1B). There were also pLMs that integrate structure token information(14, 23).

However, current protein models use biophysical information from static structures. These structural snapshots lack crucial information about thermodynamics, allostery, folding, and dynamic interactions. Moreover, static structures cannot describe the dynamic nature of IDRs, which constitute more than 30% of the human proteome(24) and use their inherent flexibility to mediate essential biological processes. Protein dynamics are fundamental to understand protein behaviors and mutation effects. Although evolutionary information encodes aspects of protein dynamics(25, 26), it depends on the quality and quantity of homologous sequences and cannot be directly extracted. Molecular dynamics (MD) simulations are routinely used to study protein dynamics, generating ensembles of structures based on defined physical forces over the simulation time. All-atom MD simulations are often computationally intensive to reach a converged equilibrium state(27). Coarse-grained models simplify protein residues into beads and use specialized force fields to reduce computation cost(24). Additionally, normal mode analysis (NMA)(28, 29) can describe protein vibrations around the equilibrium conformation, with low-frequency modes capturing large-scale motions. What’s more, deep learning models have been developed to generate conformation ensembles(30, 31). While these methods are wildly used to study protein dynamics, the produced data are often high-dimensional and irregularly shaped. Thus, effectively and efficiently integrating these dynamic data into protein deep learning models remains a significant challenge.

In this study, we present a deep learning framework to integrate dynamics data into protein language models. Using dynamics profiles from 64,403 proteins, we pre-trained (i.e., train models on large scale, general datasets to learn useful information before applying them to specific tasks) two models: SeqDance and ESMDance. SeqDance is trained from scratch directly on protein dynamics data, whereas ESMDance is built upon ESM2 outputs, which implicitly encode structural and evolutionary information. We demonstrated that our models capture local dynamic properties and global conformational properties for proteins not included in the training data. We also applied our models for zero-shot (applying models directly to a new task without any additional training) prediction of mutation effects, achieving superior performance on designed and viral proteins, where evolution-based methods typically underperform.

## Results

### Pre-training Protein Language Models on Dynamics Properties of over 64,000 Proteins

To establish the protein dynamics dataset, we curated both high-resolution and low-resolution data sources. High-resolution data includes experimental data and all-atom molecular dynamics (MD) simulation trajectories from mdCATH(32), ATLAS(33), GPCRmd(34), PED(35). Since high-resolution data is limited, we further used low-resolution dynamics data, including coarse-grained MD trajectories of human intrinsically disordered regions (IDRs) from IDRome(24) and Normal mode analysis (NMA, we used the Gaussian Network Model(28) (GNM) and the Anisotropic Network Model(29) (ANM))(28, 29) for representative structures in the PDB(36). Overall, we collected 7,799 high-resolution data and 56,604 low-resolution data, covering ordered domains, IDRs, membrane proteins, antibodies, and protein complexes (see Table 1 and SI Appendix Supplementary Methods for details).

**Table 1:** Protein dynamic datasets.

|  | Source | Description | Count | Method |
| --- | --- | --- | --- | --- |
| High-resolution | PED | Disordered regions | 382 | Experiment and others |
|  | GPCRmd | Membrane proteins | 509 | All-atom MD, ~500 ns × 3 |
|  | mdCATH | CATH domains | 5,392 | All-atom MD, ~464 ns × 5 |
|  | ATLAS | Ordered structures in PDB | 1,516 | All-atom MD, 100 ns × 3 |
| Low-resolution | IDRome | Disordered regions | 28,058 | Coarse-grained MD |
|  | Proteinflow | Ordered structures in PDB | 28,546 | Normal mode analysis |

We extracted residue-level and pairwise dynamic properties that describe the distribution of properties in structure ensembles (Figure 1C, SI Appendix Table S1). Residue-level properties include normalized root mean square fluctuation (RMSF), solvent accessible surface area (SASA), eight-class secondary structures, and dihedral angles (*phi*, *psi*, *chi1*) which describe the rotations around bonds in the protein backbone and side chains. Pairwise properties include the correlation of Cα movements and frequencies of hydrogen bonds, salt bridges, Pi-cation, Pi-stacking, T-stacking, hydrophobic interactions, and van der Waals interactions. For NMA data, we categorized normal modes of each structure into three frequency-based clusters. For each cluster, we calculated residue fluctuation and pairwise movement correlation (see SI Appendix Table S1 and Supplementary Methods for details).

We developed two models, SeqDance and ESMDance, both consisting of Transformer encoders and dynamic property prediction heads (task-specific output layers). Both models take protein sequence as the only input, use the encoder’s final layer embeddings and attention maps to predict residue-level and pairwise dynamic properties (Figure 1C). The Transformer encoder follows the same architecture as ESM2-35M (35 million parameters) with 12 layers and 20 heads per layer, the dynamic property prediction heads contain 1.2 million trainable parameters in total. The key difference between the two models lies in their initialization and use of prior knowledge. In SeqDance, all parameters were randomly initialized, allowing the model to learn dynamic properties from scratch. In contrast, ESMDance retains and freezes all parameters from ESM2-35M, enabling it to leverage the evolutionary information already encoded in ESM2 to predict dynamic properties (see Methods for details).

To balance contributions of different properties from different data sources, we adjusted their respective weights in the loss function (see Methods for details). We clustered sequences using a 50% sequence identity cutoff (same to ESM2 training), then randomly chose 95% of the clusters as the training set, reserving the remaining 5% as the test set (no validation set was used, since we did not perform hyperparameter tuning; see Methods). The training and test losses are shown in SI Appendix Figure S1. SeqDance and ESMDance achieve comparable overall test losses, with SeqDance performing better on the IDRome test set and NMA test set, while ESMDance performing better on the high-resolution test set. After pre-training, we observed strong correlations between the weights of linear layers used to predict co-movement in MD simulations and those used for co-movement in NMA, in both models (SI Appendix Figure S2). This result confirms that NMA effectively captures dynamic motions similar to those observed in MD simulations, supporting its use as a complementary source for augmenting model pre-training.

To assess the direct utility of SeqDance and ESMDance, we evaluated them on the test IDRome and high-resolution datasets, comparing performance on IDRs against the IDR conformation generative model STARLING(31) and on the high-resolution dataset against two NMA methods: GNM and ANM. We focused on residue fluctuation and pairwise movement correlation (only residue pairs farther than five amino acids were considered as recommended in CASP(37)), as they can be directly calculated from STARLING-generated ensemble and normal modes. For test IDRs, SeqDance achieves median Pearson correlations of 0.87 for residue fluctuation and 0.89 for pairwise movement correlation, ESMDance achieves median correlations of 0.82 and 0.83 for two tasks (Figure 1D–E). As a baseline, we used STARLING to generate 200 conformations for each test IDR, extracted dynamic properties from the generated ensemble. This approach achieves median correlations of 0.82 and 0.85 for two tasks (Figure 1D–E), comparable to ESMDance but not as good as SeqDance. For high-resolution ordered test proteins, SeqDance achieves median Pearson correlations of 0.54 for residue fluctuation and 0.27 for pairwise movement correlation, while ESMDance achieves 0.60 and 0.37, respectively (Figure 1F–G). As baselines, we used GNM and ANM (low-frequency 5 or 20 modes for GNM, 5×3 or 20×3 modes for ANM) to calculate dynamic properties. ESMDance is comparable to ANM in predicting residue fluctuation and comparable to GNM with five modes in predicting pairwise correlation, but overall, both SeqDance and ESMDance lag behind GNM and ANM when using 20 normal modes for ordered proteins (Figure 1F–G).

Taken together, by being trained on protein dynamic properties, SeqDance and ESMDance can be directly applied to predict residue-level and pairwise dynamic properties for proteins dissimilar to those in the training set. They demonstrate superior performance on IDRs for both residue-level and pairwise dynamics, and satisfactory performance on residue-level dynamics of ordered proteins, using sequence as the only input (NMA takes structure as input). SeqDance performs better on IDRs, whereas ESMDance performs better on ordered proteins, consistent with the trends observed in the test losses (SI Appendix Figure S1) and in line with the fact that evolutionary information is more informative for ordered proteins but less effective for IDRs. In the following, we first assessed SeqDance’s attention mechanism and embeddings, then compared its predicted dynamic properties with mutation effect on protein stability, and examined the advantage of ESMDance compared to ESM2 and SeqDance on predicting mutation effects.

### SeqDance’s Attentions Capture Dynamic Interactions and Co-movement

The Transformer model employs self-attention mechanism(38) to aggregate information from other tokens, with attention values representing the relationship between tokens (in protein context, amino acids). Here, we evaluated whether SeqDance’s attention mechanism captures dynamic interactions and residue co-movement. The analysis was conducted on the test dataset, which included 404 high-resolution proteins, 1,355 IDRome proteins, and 1,274 proteins with NMA. Additionally, we evaluated an independent test set comprising 29 MD trajectories from the Dynamic PDB dataset(39). All test proteins share less than 50% sequence identity with the SeqDance training sequences, allowing us to assess the model’s ability to generalize to unseen proteins. The Dynamic PDB test set was generated using a different MD force field to those used in pre-training, enabling us to further evaluate SeqDance’s generalization ability across different simulation conditions. For all analyses, only residue pairs farther than five amino acids were considered as recommended in CASP(37).

We compared the dynamic properties (interaction frequency and movement correlation) of residue pairs to attention values from each attention head. We calculated Spearman correlations for all 240 attention heads and highlighted the top five heads in Figure 2. For interaction frequency, the best attention heads of SeqDance achieve median Spearman correlations of 0.41 on the high-resolution test set, 0.52 on the IDRome test set, and 0.52 on the Dynamic PDB test set (Figure 2A-C; the best attention head for different data may be different). For residue pairs with positive movement correlation in the MD ensemble, SeqDance performs particularly well on the IDRome test set, achieving a median Spearman correlation of 0.74 for the best attention head. In contrast, median Spearman correlations of best attention heads are just 0.22 for both ordered test set (high-resolution and Dynamic PDB; Figure 2D-F), consistent with our results in Figure 1E, G. SeqDance’s attention do not capture residue pairs with negative movement correlation in the MD ensemble well (SI Appendix Figure S3A-C). For NMA data, the best attention heads achieve median Spearman correlations of 0.34, 0.29, and 0.22 for positively correlated pairs in low-, medium-, and high-frequency modes, respectively (Figure 2G-I), and Spearman correlations of –0.04, –0.26, and –0.35 for negatively correlated pairs in three frequency modes (SI Appendix Figure S3D-F; negative correlation means better capturing this property).

**Figure 2.**
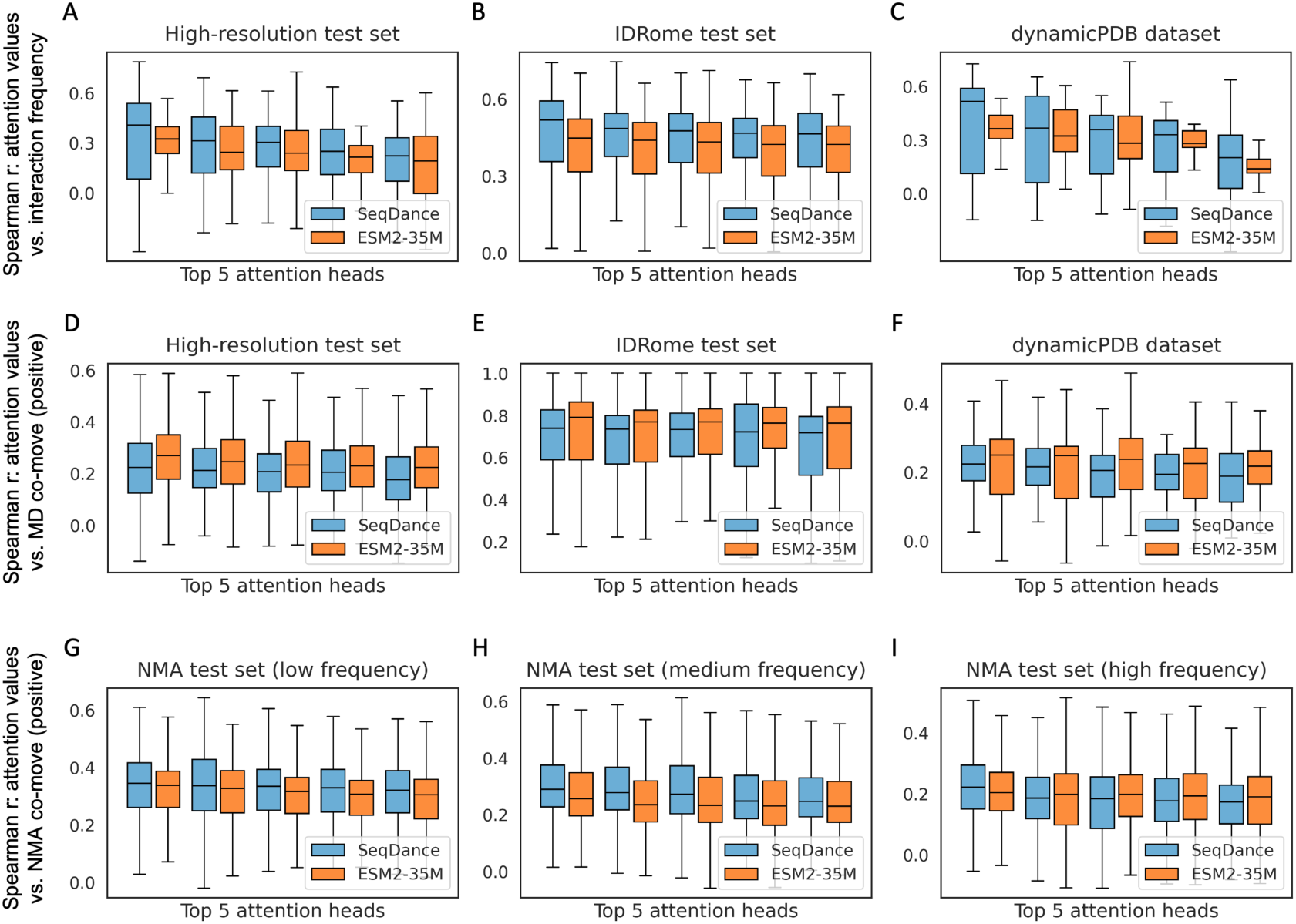
SeqDance’s attention mechanism captures dynamic interactions and co-movement in test sets. For each attention head, Spearman correlation was calculated between attention values and pairwise dynamic properties, top five heads of SeqDance and ESM2-35M are shown. **D-I** show results for positively correlated residue pairs. The boxplots show the distributions of Spearman correlation on test proteins.

We further evaluated ESM2-35M, as ESMDance was trained with ESM’s parameters frozen and thus shares identical attentions. We note that the test proteins are dissimilar to SeqDance’s training data, but appear in, or have homologs in, the ESM2 training data. Previous studies showed that ESM’s attention heads capture both native interactions observed in PDB structures(40) and non-native interactions observed in MD simulations(25) via learning evolutionary information. Across all evaluations, ESM2-35M and SeqDance show comparable performances (Figure 2; SI Appendix Figure S3). SeqDance performs slightly better on four evaluations (interaction frequencies, negatively correlated pairs in the MD ensemble, and both positively and negatively correlated pairs in NMA), while ESM2-35M performs slightly better on one evaluation (positively correlated pairs in the MD ensemble). Overall, these results demonstrated that our pre-training strategy enables SeqDance’s attention to capture dynamic interactions and residue co-movement directly from sequence, performing comparable to ESM2 on unseen proteins and different simulation conditions. ESM2’s ability to capture dynamic properties enabled us to train ESMDance, achieving comparable test loss to SeqDance with far fewer trainable parameters (SI Appendix Figure S1).

### SeqDance’s Embeddings Encode Global Protein Conformation Properties

Next, to assess SeqDance’s generalization capability, we investigated whether SeqDance learns protein conformational properties not included in the pre-training tasks. Specifically, we considered end-to-end distance (*R*_e_), asphericity, and radius of gyration (*R*_g_), which quantify motion range, deviation from a spherical shape, and compactness, respectively. Since these properties cannot be directly obtained from the model, we trained linear regression models using its embeddings in a supervised manner. To ensure a fair comparison with other pLMs in Figure 1B (ESMDance embedding is identical to ESM2-35M), we applied principal component analysis to the mean-pooled embeddings from each method and used the top 200 components as input for linear regression.

We first evaluated these pLMs on structure ensembles of IDRs(41) generated using a different coarse-grained force field compared to SeqDance training set. After removing IDRs with over 20% sequence identity and 50% coverage to any SeqDance training sequence, 28,717 IDRs remained (see Methods for details). We used mean *R*_e_, asphericity, and *R*_g_ in the ensemble (see Methods for how we account for the intrinsic length dependence of *R*_g_ and *R*_e_). The training and test sets were split in a 6:4 ratio using a 20% sequence identity cutoff to prevent information leakage. Linear regression model was trained for each pLM, and the experiment was repeated 20 times. SeqDance significantly outperforms the randomly initialized model prior to pre-training and other pLMs (Figure 3A-C), reducing the MAEs (mean absolute errors) of the random model, ESM2-650M, METL, and ProSE by 31%, 25%, 29%, and 31% on average, respectively.

**Figure 3.**
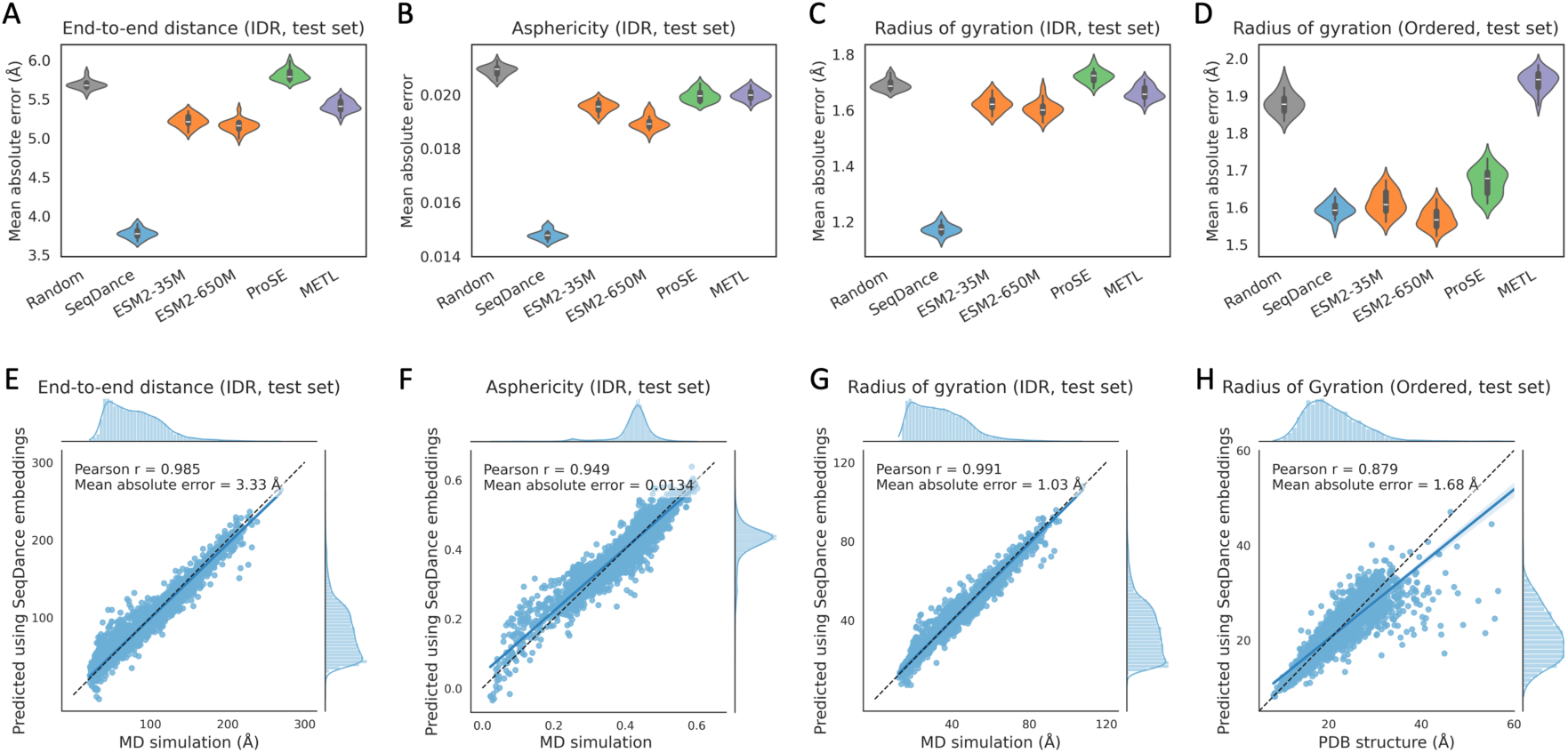
SeqDance’s embeddings encode protein global conformational properties**. A-D**. Performance comparison of SeqDance (35M parameters), METL, ProSE, and ESM2 (ESMDance embedding is identical to ESM2-35M) in predicting the end-to-end distance of disordered regions (**A**), asphericity of disordered regions (**B**), radius of gyration of disordered regions (**C**) and ordered proteins (**D**). The training and test split was 6:4 with a 20% sequence identity cutoff. Linear regression model was trained to predict conformational property using the first 200 principal components of mean-pooled embeddings from each method as input. Results presented are the distributions of the mean absolute error of 20 independent repeats. **E-G**. Comparison of conformational properties of test IDRs obtained from coarse-grained MD simulations versus predictions from SeqDance embeddings. **H**. Comparison of radius of gyration of test ordered proteins obtained from PDB structure versus predictions from SeqDance embeddings. Results in **E-H** are predictions from linear regression models using the full embeddings rather than the first 200 principal components. Each dot represents a test protein, the blue line indicates the linear fit, and the dashed black line represents the line of equality, marginal histogram shows the distribution of data.

For ordered proteins, obtaining conformational properties from structure ensembles is challenging due to the computational cost of MD simulations. Therefore, we used *R*_g_ of over 10,000 static monomer structures in the PDB database from the paper(42) (see Methods for details). Since SeqDance was trained on NMA of nearly all representative PDB structures(36), we did not exclude sequences with homologs in SeqDance training set. Using the same evaluation method described above, SeqDance outperforms the random model, ProSE, and METL, and performs comparably to ESM2 (Figure 3D).

We also evaluated the performance using Pearson correlation and found similar results (SI Appendix Figure S4A-D). Notably, even using the full embeddings instead of the first 200 principal components, SeqDance still outperforms other pLMs in most tasks (SI Appendix Table S2), despite having the shortest embedding size. Figure 3E–H shows the predictions of the linear model using full SeqDance embeddings on test proteins from one experiment. SeqDance enables accurate predictions of conformational properties observed in MD simulations of IDRs and in PDB structures of ordered proteins. We further compared SeqDance with STARLING for IDRs and with ESMFold (3B parameters) for ordered proteins. We note that both STARLING and ESMFold were trained on the same datasets used in our evaluation, meaning that a substantial fraction of the test proteins was included in their training sets. For IDRs, SeqDance outperforms STARLING when generating 20 conformations but underperforms STARLING when generating 200 conformations (Figure 3E–G; SI Appendix Figure S4E–G). For ordered proteins, SeqDance embeddings and ESMFold yield comparable performance (Figure 3H; SI Appendix Figure S4H). Thus, SeqDance provides an efficient alternative for predicting protein conformational properties, achieving performance comparable to structure- and ensemble-based methods while being orders of magnitude faster.

### SeqDance Predicted Dynamic Property Changes Reflect Mutation Effect on Protein Stability

To evaluate whether SeqDance captures information about the biophysics of protein folding, we compared its output with mutation effects on protein folding stability. We hypothesized that destabilizing mutations would induce significant changes in protein dynamics, whereas mutations with minimal impact would induce smaller dynamic changes. To test this hypothesis, we used SeqDance to predict dynamic properties for both wild-type and mutated sequences, computed their relative changes, and compared these to experimentally measured ΔΔG values (Figure 4A, see Methods for details). This approach was framed as a zero-shot prediction, as the model was not specifically trained on mutation data. We applied this approach to the mega-scale protein folding stability dataset(43), which includes ΔΔG values for 379,865 single mutations and 147,926 double mutations across 412 proteins, encompassing both natural and designed proteins.

**Figure 4.**
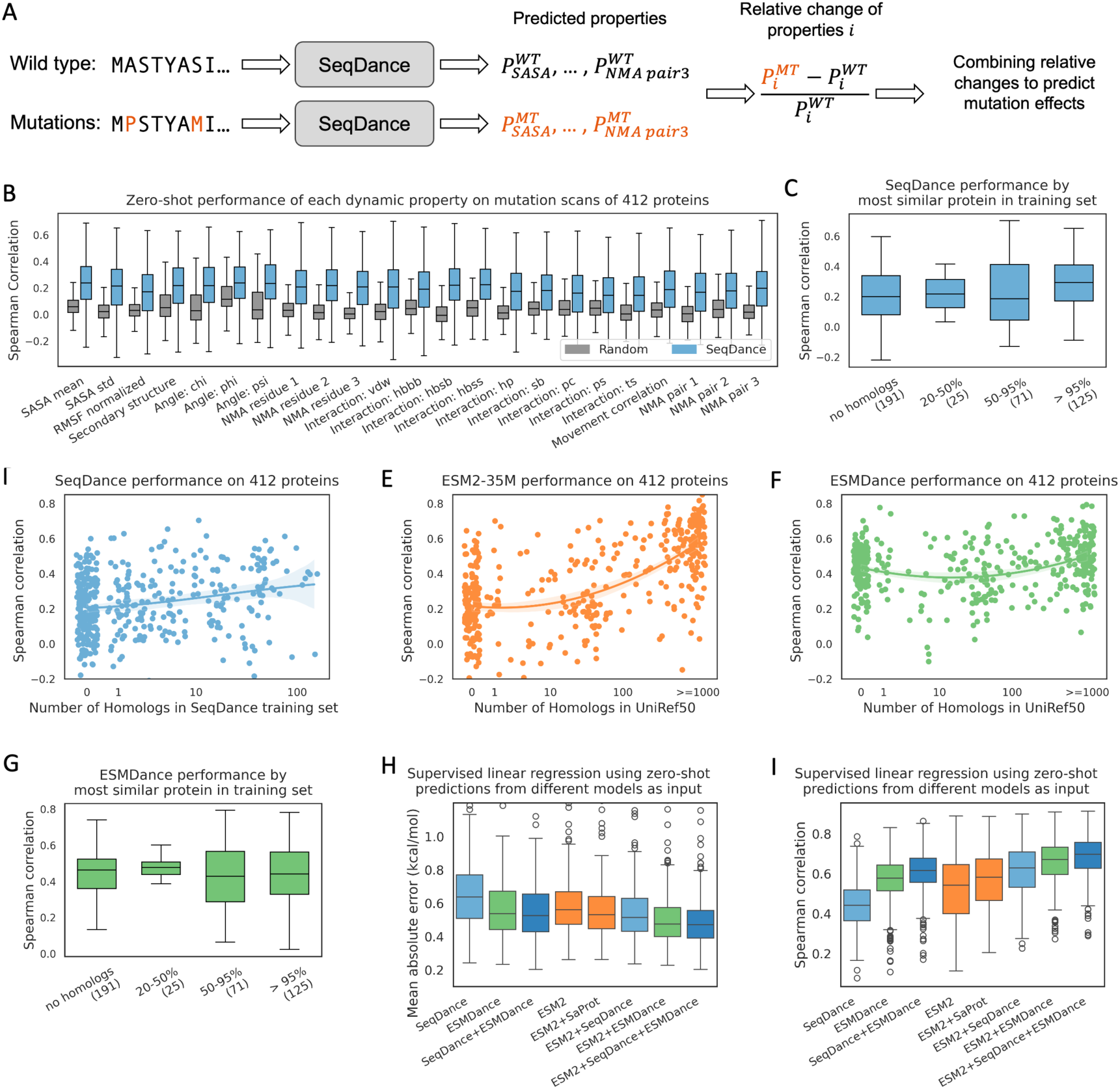
Predicting mutation effect on protein folding stability. **A.** Framework of using SeqDance (35M parameters) or ESMDance (35M parameters) to predict mutation effects. **B**. Distribution of zero-shot prediction performance (Spearman correlation) on 412 proteins of different dynamic properties; the random model (color grey) represents the randomly initialized model prior to pre-training. SASA mean, std: mean and standard deviation of solvent-accessible surface area. NMA properties 1, 2, 3: properties calculated from low-, median-, and high-frequency normal modes. vdw: van der Waals interaction; hbbb: backbone-to-backbone hydrogen bond; hbsb: side-chain-to-backbone hydrogen bond; hbss: side-chain-to-side-chain hydrogen bond; hp: hydrophobic interaction; sb: salt bridge; pc: Pi-cation interaction; ps: Pi-stacking interaction; ts: T-stacking interaction. **C**. SeqDance zero-shot prediction performance split by the most similar proteins in SeqDance training set. We used cutoff of 95% sequence identity together with 95% coverage, 50% sequence identity together with 80% coverage, 20% sequence identity together with 50% coverage. **D-F**. Relationship between zero-shot performance and the number of homologs (defined as 20% sequence identity and 50% coverage) in training set. The line represents a polynomial regression of order two, with the shaded area indicating the 95% confidence interval. The x-axis shows the log-scaled number of homologs, with small random noises added to the x values to reduce overlap. **G**. ESMDance zero-shot prediction performance split by the most similar proteins in ESMDance training set. **H-I**. Performance of supervised linear regression models on test mutations from 412 proteins. For each protein, 50% of the mutations were used to train the regression model, and the remaining 50% were reserved as the test set. For SeqDance and ESMDance, the regression models used their 23 predicted dynamic property changes as input; for ESM2, the regression model used predictions from three models (35M, 650M, and 15B) as input; and for SaProt, the zero-shot predictions of SaProt-35M were used as input.

To evaluate the zero-shot predictive power of individual dynamic properties, we computed the Spearman correlation between the relative change of each property and the corresponding ΔΔG values for mutations in each protein. While the randomly initialized model prior to pre-training shows no predictive power, many SeqDance-predicted properties achieve median Spearman correlations above 0.2 (Figure 4B; SI Appendix Figure S5), confirming that pre-training enables SeqDance to capture meaningful biophysical dynamics.

Among individual properties, solvent-accessible surface area (SASA mean) emerges as one of the top-performing properties, consistent with the fact that destabilizing mutations disrupt core packing and increase solvent exposure. Dihedral angles *phi* and *psi* also rank highly, highlighting the role of backbone conformational flexibility in protein stability. Additionally, pairwise movement correlations derived from MD simulations and NMA demonstrate relatively high performance, further emphasizing the importance of collective dynamics in predicting mutation effects. To integrate these dynamic properties, we used the geometric mean of quantile-normalized relative changes (see SI Appendix Table S3 and Methods for details). The unified score yields a median Spearman correlation of 0.24 across all 412 proteins.

To assess whether SeqDance’s performance depends on the presence of homologs in its training set, we first analyzed its performance relative to the most similar protein in the training set. SeqDance performs better on proteins with close homologs of over 95% sequence identity in its training set (Figure 4C). For the 125 proteins with such close homologs, the median Spearman correlation is 0.29, whereas for the 191 proteins with no homologs in the training set, the median correlation is 0.20. Proteins with homologs of 20–95% sequence identity show similar performance to those with no homologs. We also examined SeqDance’s performance as a function of the number of homologs (defined as over 20% sequence identity and 50% coverage) and observed only a minor correlation (Figure 4D).

Furthermore, we compared SeqDance with ESM2, which can perform zero-shot prediction of mutation effects by estimating the probability of 20 amino acids based on evolutionary information. ESM2-35M achieves a median Spearman correlation of 0.33, outperforming SeqDance. This is consistent with the fact that evolutionary information is highly effective in predicting mutation effects. However, the performance of ESM2 is strongly dependent on the number of homologs in its training set, UniRef50, particularly for proteins with over 100 homologs (Figure 4E; ESM2-650M and −15B in SI Appendix Figure S6A, B). To isolate the effect of homolog abundance, we evaluated a subset of 257 proteins that have a similar number of homologs (between 0 and 146) in both UniRef50 and the SeqDance training set. In this subset, ESM2-35M achieves a median Spearman correlation of 0.22, comparable to SeqDance’s performance of 0.24 (SI Appendix Figure S6C). These results suggest that, given similar model size and training data abundance, SeqDance and ESM2 perform similarly in predicting mutation effect on protein stability. Overall, despite being pre-trained exclusively on wild-type sequences and without reliance on evolutionary information, SeqDance is able to make meaningful predictions about mutation effect on protein stability, even for proteins with no homologs in the training set.

### ESMDance Enhances Prediction of Mutation Effect on Protein Stability

While both ESM2-35M, which leverages evolutionary information, and SeqDance, which captures protein dynamic properties, can predict mutation effect on protein stability, their individual performance remains modest at a model size of 35 million parameters. To harness the strengths of both approaches, we developed ESMDance that integrates evolutionary and dynamic information, also with 35 million parameters. Using the zero-shot approach illustrated in Figure 4A, ESMDance achieves a median Spearman correlation of 0.46 across 412 proteins (Figure 4F), significantly outperforming both ESM2-35M (0.33) and SeqDance (0.24). Remarkably, ESMDance also matches or surpasses SaProt-35M(23) (0.45; SI Appendix Figure S6D), which has similar model architecture but uses structure tokens as additional input, and much larger language models, including ESM2-650M (0.46; SI Appendix Figure S6A) and ESM2-15B (0.43; SI Appendix Figure S6B). Moreover, ESMDance demonstrates robust generalization: its performance is independent of the presence of homologs in its training set or in UniRef50 (Figure 4F-G; SI Appendix Figure S6E). We note that proteins without homologs in UniRef50 also lack homologs in the ESMDance training set. Thus, the performance gains observed for these proteins are not attributable to the presence of their dynamic data in the ESMDance training.

These models’ zero-shot predictions are correlated with ΔΔG, but not on the same physical scale. To align them with ΔΔG, we fitted supervised linear regression models. For each protein, 50% of mutations were randomly sampled for training and the remaining 50% were used for testing. Linear regression models of SeqDance- and ESMDance-predicted dynamic property changes (23 relative changes, Figure 4B) achieve median MAEs of 0.63 and 0.54 kcal/mol and median Spearman correlations of 0.44 and 0.58, respectively, on test mutations from 412 proteins (Figure 4H-I), significantly outperforming their respective zero-shot correlations. As a baseline, a linear regression model using zero-shot predictions from three ESM2 models (35M, 650M, and 15B) achieves a median MAE of 0.56 kcal/mol, better than SeqDance but worse than ESMDance (Figure 4H-I). We further trained regression models using predictions from multiple methods as inputs. Adding SaProt-35M predictions to ESM2 decreases the MAE to 0.53 kcal/mol, while adding SeqDance or ESMDance predictions yields larger improvements: ESM2 + SeqDance reaches 0.52 kcal/mol, ESM2 + ESMDance reaches 0.48 kcal/mol, and combining ESM2, SeqDance, and ESMDance achieves the best performance with a median MAE of 0.47 kcal/mol and a median Spearman correlation of 0.70 (Figure 4H–I).

Overall, these results demonstrate that by integrating evolutionary information learned by ESM2 with protein dynamic properties, ESMDance achieves superior performance in predicting mutation effect on protein stability. Furthermore, our models provide additional information to evolution-based models, and their combination yields even greater predictive power.

### Biophysics-based Pre-training Improves Mutation Effect Prediction for Designed and Viral Proteins

One limitation of ESM2 is that the evolutionary information it learns from natural proteins may fail to generalize to de novo designed proteins. To evaluate whether ESMDance can address this limitation, we analyzed a subset of 135 designed proteins that lack homologs in both UniRef50 and the SeqDance/ESMDance training set. ESM2-35M, −650M, and −15B achieve median Spearman correlations of 0.21, 0.32, and 0.29, respectively—significantly lower than their average performances (Figure 5A, 4E; SI Appendix Figure S6A, S6B). In contrast, ESMDance achieves a median Spearman correlation of 0.46, substantially outperforming SeqDance (0.20), SaProt-35M (0.06) and all ESM2 models (Figure 5A). Further analysis at the individual protein level revealed that the performances of different ESM2 models are highly correlated (Figure 5B). In contrast, both SeqDance and ESMDance exhibit orthogonal performance patterns to ESM2 (Figure 5C-D), with ESMDance showing significant improvements for most proteins compared to ESM2-35M (Figure 5D).

**Figure 5.**
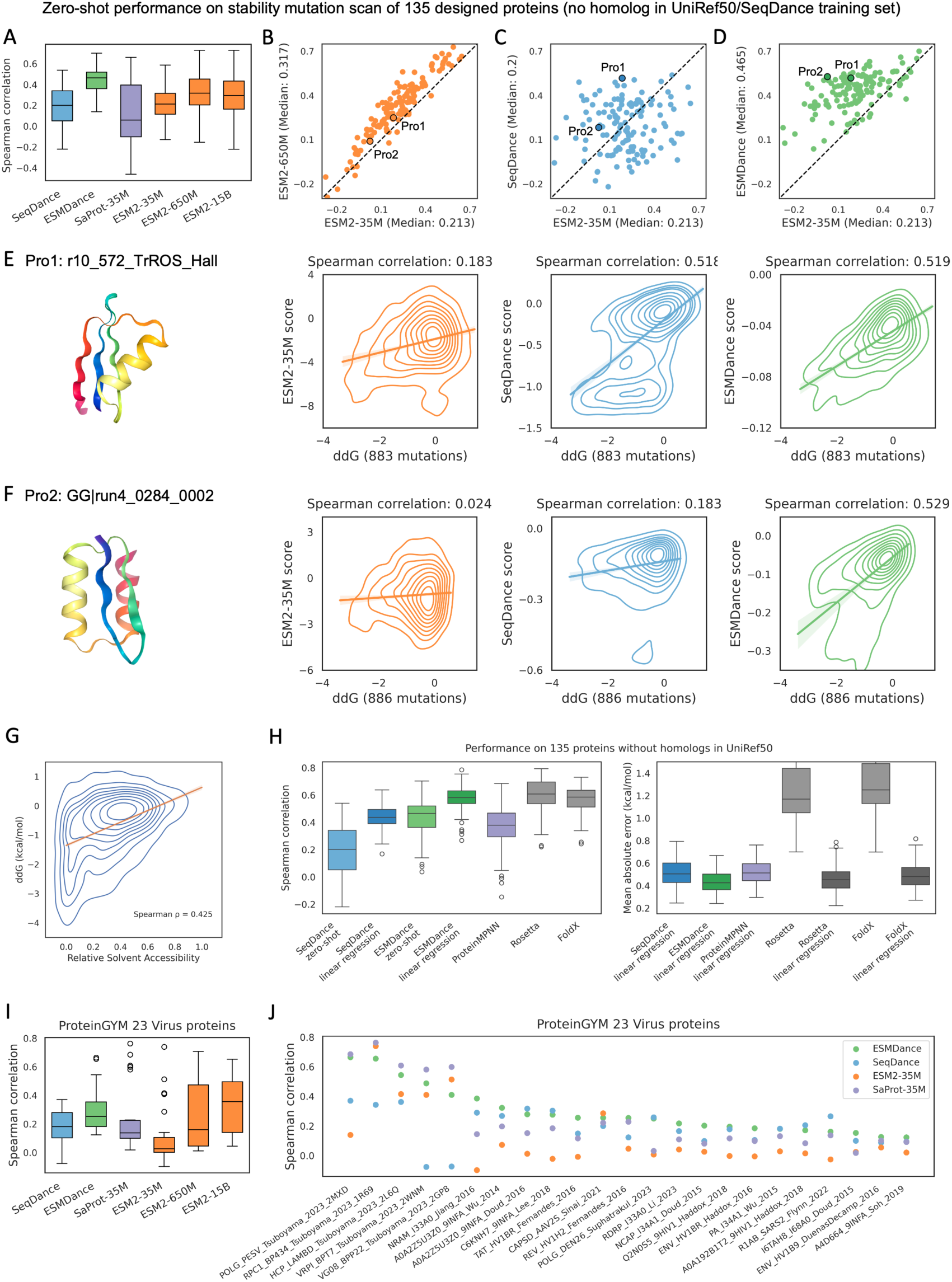
Predicting mutation effects for designed and viral proteins. **A-D.** Performance comparison between ESM2, SaProt (35M parameters), SeqDance (35M parameters), and ESMDance (35M parameters) for 135 designed proteins with no homologs in both UniRef50 and the SeqDance training set. Two highlighted proteins are further shown in E-F. **E-F.** Structure and the relationship between predictions and ΔΔG values for two designed proteins. The kernel density estimate plot shows the distribution of experimentally measured folding ΔΔG values and zero-shot prediction values for the three methods. The line represents the linear regression, with the shaded area indicating the 95% confidence interval. **G**. Relationship between experimental ΔΔG and relative solvent-accessible surface area of the mutation site. The line represents the linear regression fit. **H**. The SeqDance/ESMDance zero-shot prediction uses the geometric mean of 23 relative changes in dynamic properties, whereas the SeqDance/ESMDance linear-regression model uses a linear combination of the same 23 relative changes. For all linear-regression models, 50% of mutations were randomly sampled as training set for each protein, and the remaining 50% were used as test, the performances on test mutations were shown here. **I-J.** Zero-shot performance of ESM2, SaProt, SeqDance, and ESMDance on 23 viral proteins shorter than 1024 residues in ProteinGYM.

Furthermore, we visualized zero-shot predictions for two designed proteins (highlighted in Figure 5B-D). The first, r10_572_TrROS_Hall (Figure 5E), was designed using TrRosetta hallucination(43). Both SeqDance and ESMDance achieve significantly higher correlations (0.52 for both) compared to the ESM2 models (0.18, 0.25, and 0.20 for ESM2-35M, −650M, and −15B, respectively). The second, GG|run4_0284_0002 (Figure 5F), was designed using the EEHH method(43). Here, both ESM2 and SeqDance perform poorly (0.02, 0.09, −0.02, and 0.18 for ESM2-35M, −650M, −15B, and SeqDance, respectively), whereas ESMDance achieves a markedly higher correlation of 0.53. Linear regression models show similar results on test mutations of two proteins (SI Appendix Figure S7). This result underscores the advantage of ESMDance in directly integrating dynamic properties and evolutionary information during model training, outperforming separate predictions from two models.

Additionally, we benchmarked several established structure-based models. These included the inverse-folding model ProteinMPNN(44), which predicts sequence from structure and whose predicted sequence likelihood correlates with mutation effects on stability, as well as the biophysics-based models Rosetta(45) and FoldX(46) (see Methods for details), which directly estimate mutation ΔΔG values. We note that using the structure as input inherently simplifies stability prediction, since mutations in buried sites typically have larger effects, whereas surface mutations tend to have smaller effects (Figure 5G). We first evaluated Spearman correlation, ESMDance zero-shot outperforms ProteinMPNN but performs below FoldX and Rosetta, while ESMDance linear-regression performs comparably to FoldX and Rosetta (Figure 5H). We also assessed the mean absolute error (MAE), and our linear-regression models yield substantially lower MAE than both FoldX and Rosetta raw ΔΔG values. After calibrating FoldX and Rosetta raw ΔΔG values with experimental measurements using linear regression, their MAE becomes comparable to—but still slightly higher than—that of ESMDance linear-regression model (Figure 5H). Overall, these results suggest that our linear-regression models can serve as an alternative to traditional biophysics-based models while being orders of magnitude faster and requiring only sequence input.

Besides designed proteins, we also examined their performances on viral proteins, which evolve rapidly and are underrepresented in UniProt. For deep mutational scanning experiments in ProteinGYM(47), ESM2 models and SaProt-35M consistently perform worse on viral proteins compared to the other proteins regardless of model size (SI Appendix Figure S8). ESM2-35M achieves a median Spearman correlation around 0 on viral proteins (SI Appendix Figure S8). For 23 viral proteins shorter than 1024 residues, encompassing 187,902 mutations, SeqDance and ESMDance achieve median Spearman correlations of 0.18 and 0.25, respectively (Figure 5I-J), using the zero-shot approach in Figure 4A (which may not be the optimal strategy for mutational scanning beyond stability, see Discussion). They outperform ESM2-35M (0.03) and SaProt-35M (0.14) but are not as good as ESM2-15B (0.36; Figure 5I).

## Discussion

In this work, we developed SeqDance and ESMDance, two protein language models (pLMs) pre-trained with dynamic properties derived from molecular dynamics (MD) simulations and normal mode analysis (NMA). SeqDance captures local dynamic interactions, residue co-movement, and global conformational properties after pre-training, using only single sequence input and without relying on evolutionary information. ESMDance integrates the evolutionary information encoded in ESM2 and the newly learned dynamic properties, significantly outperforms SeqDance and ESM2-35M in zero-shot prediction of mutation effects for designed and viral proteins.

In the pre-training process, we did not directly predict full structure ensembles due to the immense size of the dataset (over 60 million frames) and the complexity of modeling entire ensembles. Instead, we used simplified dynamic property descriptors such as the mean, standard deviation, and interval distribution of ensemble-derived properties. Prior study has demonstrated that mean values from structure ensembles provide significantly more information than those from static structures(48). This strategy enables SeqDance and ESMDance to learn from simplified but informative properties of protein dynamics, without the overwhelming computational demand of full ensemble modeling. During pretraining, the test losses plateaued early (SI Appendix Figure S1), but we continued training to enable models to learn dynamic information more effectively from the training set, as the training set was representative and relatively large, and extended training did not result in increased test loss. Using attention maps to predict pairwise properties constrains SeqDance to focus on interacting and co-moving residue pairs. This approach is crucial for learning biophysically meaningful information and improves SeqDance’s ability to extrapolate to unseen proteins. Our results show that our pre-training strategy enables the transformer’s attentions to effectively capture pairwise dynamic properties, using only 1/1000th of the training sequences compared to ESM2.

In Figures 1–2, both SeqDance and ESMDance achieve higher correlations for IDRs compared to ordered structures, likely because our models were trained on more IDR MD trajectories. In addition, the dynamics of IDRs are usually localized and easier to learn, whereas the dynamics of ordered structures often involve long-range interactions. SeqDance effectively learns global conformational properties of both ordered and disordered proteins (Figure 3), which are essential for understanding protein shape and flexibility. In comparison, METL(22) underperforms in predicting the radius of gyration (*R*_e_) for ordered proteins (Figure 3D), despite having *R*_g_ prediction as a pre-training task. ProSE(9) performs well in predicting the conformational properties of ordered regions but struggles with IDRs (Figure 3), likely because its pre-training focused on ordered PDB structures. ESM2 performs well on both ordered and disordered proteins, as conformational properties are also conserved(41).

Current methods for zero-shot prediction of mutation effects use evolutionary information and static structure. These methods are less effective for proteins without sufficient homologs and structures. We applied SeqDance and ESMDance to predict mutation effects by calculating changes of dynamic properties between wild-type and mutated sequences. These results highlight the effectiveness and efficiency of integrating both evolutionary information and dynamic properties for understanding mutation effects, particularly for proteins that cannot be accurately predicted using evolutionary information or static structure. While, to our knowledge, we are the first to apply this kind of zero-shot strategy to predict mutation effects on protein, analogous approaches like Enformer(49) and Borzoi(50) have been used to predict effect of non-coding mutations in DNA sequences, where evolutionary information is also less effective. Our current zero-shot strategy uses changes across the entire protein, which is reasonable for predicting mutation effects on protein stability but not optimal for mutation effects on function, binding, or activity, where dynamic changes at functional sites, binding interfaces, or catalytic centers may be more relevant. Moreover, for SeqDance, the presence of highly similar proteins in the training set leads to improved performance on mutation effect (Figure 4C). Thus, running MD simulations for a protein of interest and using the extracted features to fine-tune SeqDance may yield better predictions, but this may not work for ESMDance (Figure 4G).

We did not compare our models to classical protein dynamics methods on specific proteins. This was a deliberate choice: our models are not intended as drop-in replacements for existing approaches, but rather as a first step toward a new paradigm that learn dynamic information with protein language models. In our work, all evaluations were performed on proteins shorter than 1,024 residues. We envision several directions to further improve the models. First, expanding protein dynamics data: while high-resolution data remain limited relative to the vast number of sequences, lower-resolution sources, such as coarse-grained MD simulations and NMA, can still provide valuable training signals. Second, training models on more detailed dynamic properties that describe higher-order relationships and time-dependent behaviors to capture more nuanced aspects of protein motion. Third, employing more advanced model architectures (e.g., the Evoformer in AlphaFold2(1)) and scaling up model size: as demonstrated in many areas of deep learning, larger models can learn more complex patterns. Lastly, incorporating structural information as additional input may further improve performance on ordered structures and mutation effect on stability.

In conclusion, SeqDance and ESMDance advance the integration of protein dynamic properties, captured through molecular dynamics simulations and normal mode analysis, into deep learning frameworks. By complementing widely used evolutionary and static structure-based models, our models reveal novel insights into protein behavior and mutation effects, particularly for proteins lacking homologs, such as IDRs, designed and viral proteins. This underscores the practical utility of dynamic properties as a critical, yet underexplored, dimension in computational protein design and studies, offering a new tool to bridge the gap between sequence, dynamics, and function.

## Methods

### Model Architecture

The models consist of a transformer encoder and the dynamic property prediction heads. The transformer encoder uses the ESM2-35M architecture and the same sequence tokenizer. It comprises 12 layers with 20 attention heads each and employs rotational positional embeddings, with a final embedding dimension of 480. The choice of the 35M model is due to both the relatively small training dataset (compared to hundreds of millions of sequences) and our computational resources. Linear layers are added to predict residue-level properties from the residue embeddings of the last transformer layer. For pairwise feature prediction, pairwise embeddings are computed from residue embeddings as follows:

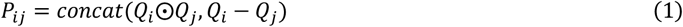

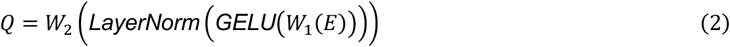

where *E* represents the final residue embeddings, *W*_1_ and *W*_2_ are learnable linear matrices. *i* and *j* are residue indexes, ⊙ represents element-wise multiplication, The pairwise embeddings *P*_ij_, along with attention values from 240 heads, are passed through a linear layer to predict pairwise features. The dynamic property prediction heads contain 1.2 million parameters in total.

Using attention maps and pairwise embeddings allows the model to capture the distinct characteristics of two pairwise properties: the mean value of interaction frequency depends on the sequence length, as the maximum number of interactions a residue can form is constrained, while the mean value of movement correlation is length-independent. By combining length-dependent attention values (which sum to one after SoftMax operation in attention calculation) with length-independent pairwise embeddings, the model can capture both types of pairwise properties.

Different activation functions were used for different properties: For secondary structure and dihedral angles, where the values for each property sum to one, SoftMax functions were used. For RMSF, which is normalized to (0,1], Sigmoid functions were applied. For correlation properties, Tanh functions were used to maintain values within (−1,1). For other non-negative properties, SoftPlus functions were applied (ReLU led to the dead ReLU problem in our experiments).

### Pre-training Procedure

Models were implemented and trained using PyTorch(51) (v2.5.1) and the Transformers library (v4.48.2, https://huggingface.co/docs/transformers/index). Training was performed with a batch size of 128, parameters were updated every three batches, each from high-resolution data, IDRome, and the NMA dataset. Optimization was conducted using the AdamW optimizer, with a peak learning rate of 1×10⁻⁴, an epsilon value of 1×10⁻⁸, betas of (0.9, 0.98), and a weight decay of 0.01. The learning rate was linearly warmed up over the first 2,000 steps, followed by a linear decay to 1×10⁻⁵ over 90% of total training steps. To mitigate overfitting, a dropout rate of 0.1 was applied in both the transformer encoder and the dynamic property prediction heads.

SeqDance was trained for 200,000 steps, with a maximum sequence length of 256 for the first 160,000 steps and 1024 for the remaining 40,000 steps. ESMDance was trained for 60,000 steps, using a maximum sequence length of 256 for the first 40,000 steps and 1024 for the final 20,000 steps. ESMDance has fewer trainable parameters, both its training and test losses plateaued earlier (SI Appendix Figure S1). For sequences exceeding the maximum length, a region of the max length was randomly selected, the selected regions were different in each epoch. Two models were trained for ten days using four Nvidia L40s GPUs. Due to computational constraints, hyperparameter tuning was not performed, which means the current model configuration may not be the best.

### Adjusting Weights of Dynamic Properties in the Loss Function

Root mean squared error (RMSE) was used as the loss function for pre-training. For interaction frequencies, some interacting pairs exhibit frequencies close to zero, which are meaningful but treated similarly to zero in RMSE. To address this, we applied a cube root transformation to scale up low-frequency interactions.

The models were pre-trained on multiple tasks of varying scales. To balance the losses across tasks, we used the standard deviations of dynamic properties to calculate task-specific weights. The standard deviation of each property for each protein in the training set was computed. For multi-class features, such as secondary structures, the losses were calculated for each class and then averaged. These values were used to adjust the task-specific weights. Given the higher complexity of pairwise properties compared to residue-level properties, we set the pairwise-to-residue loss ratio to 3:1. For NMA data, this ratio was adjusted independently for the three frequency ranges. The adjusted losses for SASA, RMSF, secondary structure, and dihedral angles were kept equal. The adjusted losses for the nine interactions were kept equal. The ratio between the summed interaction loss and the co-movement loss was set to 3:1. The final losses for high-resolution data, IDRome data, and NMA data were further adjusted to be equal. The final weights for each property are available at https://huggingface.co/datasets/ChaoHou/protein_dynamic_properties.

### Attention analysis

The attention map and the interaction map are not symmetric (the interactions have direction), therefore, the means of the original matrixes and its transposes were used. Only residues farther than five amino acids in protein sequence are considered. The interaction frequencies of all nine interaction types were summed for analysis. Only proteins shorter than 1,024 residues were analyzed.

### Evaluation of Conformational Properties for IDRs and Ordered Structures

Conformational properties asphericity, end-to-end distance (*R*_e_) and radius of gyration (*R*_g_) from coarse-grained MD trajectories of over 40,000 IDRs(41) were downloaded from https://zenodo.org/records/10198621. Only IDRs ranging from 32 to 384 residues in length were included (the sequence cutoff was used because we compared our model with STARLING, which can only generate conformation for sequence no longer than 384), IDRs with over 20% sequence identity and 50% coverage (determined using MMseqs2 search with parameters: *-s 7 -a 1*) to any SeqDance training sequence were removed, resulting in a final set of 28,717 IDRs. *R*_g_ values were calculated for monomer PDB structures analyzed in the paper(42) using mdtraj, only proteins no longer than 800 were analyzed (the sequence cutoff was used because we compared our model with ESMFold, which runs out of GPU memory when processing long proteins).

Both *R*_e_ and *R*_g_ were first normalized to account for their inherent dependence on sequence length, as follows:

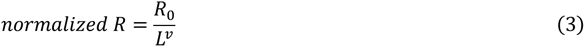

Where *L* is the sequence length, and *R*_e_ is a prefactor determined by the biochemical properties of the protein, ν is the scaling exponent. The ν values for both *R*_e_ and *R*_g_ (for ordered protein and IDR separately) were obtained by fitting a linear regression model between *log*(*R*) and *log*(*L*).

For both IDRs and ordered structures, MMseqs2 easy-cluster was used to cluster sequences with the parameters: *--min-seq-id 0.2 -c 0.5 --cov-mode 0*. Sequences from 60% of randomly selected clusters were used as the training set, while the remaining sequences formed the test set. SeqDance was compared with METL (METL-G-50M-1D), ESM2 models, and ProSE. The last layer embeddings of SeqDance, METL, and ESM2, and concatenated embeddings of all layers of ProSE (as suggested in their GitHub: https://github.com/tbepler/prose) were used. All embeddings were mean-pooled along protein sequences. Linear regression (scikit-learn v1.4.1) with default parameters was used to predict conformational properties (asphericity, normalized *R*_e_, and normalized *R*_g_), trained separately for the embeddings from each method. The predicted normalized *R*_e_ and normalized *R*_g_ were then multiplied with *L*^2^ to get the final prediction. Principle component analysis was conducted with scikit-learn.

STARLING was used to generate structure ensembles of IDRs with default parameters. ESMFold (3 billion parameters) was used to predict structure of ordered protein.

### Prediction of Mutation Effects

For SeqDance and ESMDance, the dynamic properties of both wild-type and mutated sequences were predicted. For property *i*, the zero-shot score *S*_i_ is computed as:

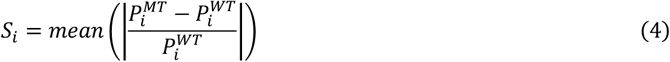

where *P*_i_^WT^ and *P*_i_^MT^ represent the predicted values of property *i* for the wild-type and mutated sequences, respectively. The relative change was computed element-wise. For example, in the case of secondary structure with eight classes, the resulting relative change matrix had a shape of *L×8* for a protein of length *L*, and the mean of absolute values of this matrix was used as the zero-shot score of secondary structure.

To combine scores from different methods, multiple aggregation strategies were evaluated, including mean, max, weighted mean, and geometric mean, applied to both raw and quantile-normalized scores. Quantile normalization ensures that different score distributions are same by aligning their values as mean of raw values of each rank. Weighted mean and geometric mean were calculated as:

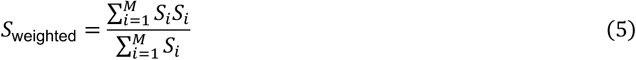

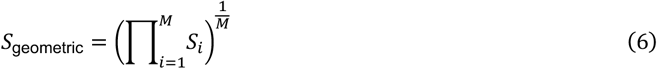

Where *M* is the number of all properties. All these zero-shot scores are non-negative. To compare them with negative ΔΔG values, the negative of the zero-shot scores was used. Various strategies to combine the relative changes of different dynamic properties were tried but not improvement over using individual properties such as mean SASA or dihedral angle *psi* was observed. Users may use these two individual properties for applications. SeqDance and ESMDance are not suitable for cross-protein prediction in our experiment, as the effects of dynamic property changes vary between proteins.

For ESM2 models and SaProt, we followed the method used in ProteinGYM. Each residue in a protein was sequentially masked, the model was used to predict the probabilities of 20 amino acids for the masked position. For a protein of length *L*, the model was run *L* times to obtain the wild-type and mutated residue probabilities at each site. The log-likelihood ratio (LLR) was then calculated for each mutation. For double mutations, the LLR scores of the two single mutations were summed to approximate the combined effect. For ESM2-15B, half-precision (float16) was used, as the model could not be loaded onto a single GPU with 48 GB of memory. For SaProt, structure tokens were used as additional input. For ProteinMPNN(44) (vanilla_model_weight v_48_020), full-sequence likelihoods were used to infer mutation effects. For FoldX(46), the structures were first repaired using the RepairPDB command, and the BuildModel command was then applied to compute ΔΔG values. For Rosetta, ΔΔG values were calculated using the RosettaDDGPrediction(45) protocol (Rosetta version 2022.11, cartddg2020_ref2015). Because Rosetta is computationally expensive, 100 mutations per protein were randomly sampled for evaluation. AlphaFold-predicted protein structures were used as input for these models.

### Mutation Datasets

For the evaluation of protein folding stability, the dataset “Tsuboyama2023_Dataset2_Dataset3_20230416.csv” and AlphaFold-predicted protein structures were downloaded from https://zenodo.org/records/7992926. Proteins were defined using the “WT_name” column in the table. Insertions and deletions were removed, as well as substitution mutations without “ddG_ML” values. The mean “ddG_ML” values for the same sequence were used. Homologs in both the SeqDance/ESMDance model training set and UniRef50 (downloaded in February 2025) were identified using MMseqs2 search with the parameters: *-s 7 -a 1 --max-seqs 1000*. The surface area was calculated using MDTraj, relative surface area was calculated by dividing the solvent-accessible surface area of each residue by the maximum surface area of the corresponding amino acid across all proteins. ProteinGYM data and predictions for these mutations were downloaded from its website (https://proteingym.org/) and GitHub in June 2025.

## Acknowledgments

This work was supported by NIH grants (R35GM149527) and Simons Foundation SFARI #1019623. We thank Dr. Guojie Zhong, Aziz Zafar, and Di Liu for helpful discussions and suggestions.

## Data availability

The extracted dynamic properties (approximately 100 GB in total), sequences (including train-test splits), loss weights for different properties, and all other data required to reproduce main results in this paper are available at: https://huggingface.co/datasets/ChaoHou/protein_dynamic_properties.

## Code availability

The codes for feature extraction, model training and evaluations are available on GitHub: https://github.com/ShenLab/SeqDance. Model weights are available at: https://doi.org/10.5281/zenodo.15047777.

## References

1. J. Jumper, et al., Highly accurate protein structure prediction with AlphaFold. Nature (2021). 10.1038/s41586-021-03819-2.

2. J. Abramson, et al., Accurate structure prediction of biomolecular interactions with AlphaFold 3. Nature (2024). 10.1038/s41586-024-07487-w.

3. M. Baek, et al., Accurate prediction of protein structures and interactions using a three-track neural network. Science 373, 871–876 (2021).

4. Z. Lin, et al., Evolutionary-scale prediction of atomic-level protein structure with a language model. Science 379, 1123–1130 (2023).

5. M. Kulmanov, et al., Protein function prediction as approximate semantic entailment. Nature Machine Intelligence 6, 220–228 (2024).

6. C. Hou, Y. Li, M. Wang, H. Wu, T. Li, Systematic prediction of degrons and E3 ubiquitin ligase binding via deep learning. BMC Biol 20, 162 (2022).

7. H. R. Kilgore, et al., Protein codes promote selective subcellular compartmentalization. Science 387, 1095–1101 (2025).

8. P. Gainza, et al., Deciphering interaction fingerprints from protein molecular surfaces using geometric deep learning. Nat Methods 17, 184–192 (2020).

9. T. Bepler, B. Berger, Learning the protein language: Evolution, structure, and function. Cell Syst 12, 654–669 e3 (2021).

10. A. Rives, et al., Biological structure and function emerge from scaling unsupervised learning to 250 million protein sequences. Proc Natl Acad Sci U S A 118 (2021).

11. A. Elnaggar, et al., ProtTrans: Toward Understanding the Language of Life Through Self-Supervised Learning. IEEE Trans Pattern Anal Mach Intell 44, 7112–7127 (2022).

12. A. Madani, et al., Large language models generate functional protein sequences across diverse families. Nat Biotechnol 41, 1099–1106 (2023).

13. Z. Zhang, et al., Protein language models learn evolutionary statistics of interacting sequence motifs. Proc Natl Acad Sci U S A 121, e2406285121 (2024).

14. T. Hayes, et al., Simulating 500 million years of evolution with a language model. Science 387, 850–858 (2025).

15. G. Zhong, Y. Zhao, D. Zhuang, W. K. Chung, Y. Shen, PreMode predicts mode-of-action of missense variants by deep graph representation learning of protein sequence and structural context. Nat Commun 16, 7189 (2025).

16. H. Zhang, M. S. Xu, X. Fan, W. K. Chung, Y. Shen, Predicting functional effect of missense variants using graph attention neural networks. Nat Mach Intell 4, 1017–1028 (2022).

17. F. Ding, J. Steinhardt, Protein language models are biased by unequal sequence sampling across the tree of life. bioRxiv 2024.03.07.584001 (2024). 10.1101/2024.03.07.584001.

18. M. Akdel, et al., A structural biology community assessment of AlphaFold2 applications. Nat Struct Mol Biol 29, 1056–1067 (2022).

19. R. M. Vernon, et al., Pi-Pi contacts are an overlooked protein feature relevant to phase separation. eLife 7, e31486 (2018).

20. H. M. Berman, et al., The Protein Data Bank. Nucleic acids research 28, 235–242 (2000).

21. C. Hou, et al., PhaSepDB in 2022: annotating phase separation-related proteins with droplet states, co-phase separation partners and other experimental information. Nucleic Acids Res 51, D460–D465 (2023).

22. S. Gelman, et al., Biophysics-based protein language models for protein engineering. Nat Methods 22, 1868–1879 (2025).

23. J. Su, et al., SaProt: Protein Language Modeling with Structure-aware Vocabulary. bioRxiv 2023.10.01.560349 (2024). 10.1101/2023.10.01.560349.

24. G. Tesei, et al., Conformational ensembles of the human intrinsically disordered proteome. Nature (2024). 10.1038/s41586-023-07004-5.

25. A. Fung, A. Koehl, M. Jagota, Y. S. Song, The Impact of Protein Dynamics on Residue-Residue Coevolution and Contact Prediction. bioRxiv 2022.10.16.512436 (2022). 10.1101/2022.10.16.512436.

26. H. K. Wayment-Steele, et al., Predicting multiple conformations via sequence clustering and AlphaFold2. Nature (2023). 10.1038/s41586-023-06832-9.

27. S. Piana, K. Lindorff-Larsen, D. E. Shaw, Atomic-level description of ubiquitin folding. Proceedings of the National Academy of Sciences 110, 5915–5920 (2013).

28. T. Haliloglu, I. Bahar, B. Erman, Gaussian Dynamics of Folded Proteins. Physical Review Letters 79, 3090–3093 (1997).

29. A. R. Atilgan, et al., Anisotropy of fluctuation dynamics of proteins with an elastic network model. Biophys J 80, 505–15 (2001).

30. S. Lewis, et al., Scalable emulation of protein equilibrium ensembles with generative deep learning. Science 0, eadv9817 (2025).

31. B. Novak, J. M. Lotthammer, R. J. Emenecker, A. S. Holehouse, Accurate predictions of conformational ensembles of disordered proteins with STARLING. [Preprint] (2025). Available at: https://www.biorxiv.org/content/10.1101/2025.02.14.638373v1 [Accessed 9 October 2025].

32. A. Mirarchi, T. Giorgino, G. De Fabritiis, mdCATH: A Large-Scale MD Dataset for Data-Driven Computational Biophysics. Sci Data 11, 1299 (2024).

33. Y. Vander Meersche, G. Cretin, A. Gheeraert, J. C. Gelly, T. Galochkina, ATLAS: protein flexibility description from atomistic molecular dynamics simulations. Nucleic Acids Res 52, D384–D392 (2024).

34. I. Rodriguez-Espigares, et al., GPCRmd uncovers the dynamics of the 3D-GPCRome. Nat Methods 17, 777–787 (2020).

35. H. Ghafouri, et al., PED in 2024: improving the community deposition of structural ensembles for intrinsically disordered proteins. Nucleic Acids Res 52, D536–D544 (2024).

36. Y. Mi, S.-B. Marcu, V. V. B. Yallapragada, S. Tabirca, ProteinFlow: An advanced framework for feature engineering in protein data analysis. Biotechnology and Bioengineering 121, 3563–3571 (2024).

37. A. Kryshtafovych, T. Schwede, M. Topf, K. Fidelis, J. Moult, Critical assessment of methods of protein structure prediction (CASP)-Round XIII. Proteins 87, 1011–1020 (2019).

38. A. Vaswani, et al., Attention is all you need in (2017), pp. 5998–6008.

39. C. Liu, et al., Dynamic PDB: A New Dataset and a SE(3) Model Extension by Integrating Dynamic Behaviors and Physical Properties in Protein Structures. [Preprint] (2024). Available at: http://arxiv.org/abs/2408.12413 [Accessed 9 October 2025].

40. R. Rao, J. Meier, T. Sercu, S. Ovchinnikov, A. Rives, Transformer protein language models are unsupervised structure learners. bioRxiv 2020.12.15.422761 (2020). 10.1101/2020.12.15.422761.

41. J. M. Lotthammer, G. M. Ginell, D. Griffith, R. J. Emenecker, A. S. Holehouse, Direct prediction of intrinsically disordered protein conformational properties from sequences. Nat Methods (2024). 10.1038/s41592-023-02159-5.

42. J. J. Tanner, Empirical power laws for the radii of gyration of protein oligomers. Acta Crystallogr D Struct Biol 72, 1119–1129 (2016).

43. K. Tsuboyama, et al., Mega-scale experimental analysis of protein folding stability in biology and design. Nature 620, 434–444 (2023).

44. J. Dauparas, et al., Robust deep learning–based protein sequence design using ProteinMPNN. Science 378, 49–56 (2022).

45. V. Sora, et al., RosettaDDGPrediction for high-throughput mutational scans: From stability to binding. Protein Sci 32, e4527 (2023).

46. J. Schymkowitz, et al., The FoldX web server: an online force field. Nucleic Acids Res 33, W382–8 (2005).

47. P. Notin, et al., ProteinGym: Large-Scale Benchmarks for Protein Fitness Prediction and Design. Advances in Neural Information Processing Systems 36, 64331–64379 (2023).

48. S. Gu, et al., Can molecular dynamics simulations improve predictions of protein-ligand binding affinity with machine learning? Brief Bioinform 24 (2023).

49. Ž. Avsec, et al., Effective gene expression prediction from sequence by integrating long-range interactions. Nature Methods 18, 1196–1203 (2021).

50. J. Linder, D. Srivastava, H. Yuan, V. Agarwal, D. R. Kelley, Predicting RNA-seq coverage from DNA sequence as a unifying model of gene regulation. Nature Genetics (2025). 10.1038/s41588-024-02053-6.

51. A. Paszke, et al., PyTorch: An Imperative Style, High-Performance Deep Learning Library. [Preprint] (2019). Available at: http://arxiv.org/abs/1912.01703 [Accessed 9 October 2025].

