## Supplementary materials for "Protein Language Models Trained on Biophysical Dynamics Inform Mutation Effects"

**This PDF file includes:**

Supporting text

Figures S1 to S8

Tables S1 to S3

SI References

### Supporting Information Text

### Supplementary Methods

#### Protein Dynamic Data Collection and Processing

The Protein Ensemble Database[1] (PED) provides conformational ensembles for intrinsically disordered proteins, primarily derived from experiments, with some from molecular dynamics (MD) simulations or predictions. All available ensembles in PED were downloaded, and sequences shorter than 16 residues were excluded. GPCRmd[2] is a community-driven database of MD simulations of G-protein-coupled receptors (GPCRs), with most proteins simulated with the CHARMM force field for 500 ns in three replicates. mdCATH[3] provides MD simulations for 5,398 CATH domains ranging in length from 50 to 500 residues, using the CHARMM22 force field. Each domain was simulated at five temperatures (320 K, 348 K, 379 K, 413 K, and 450 K), with five replicas per temperature, averaging 464 ns per replica. Five replicas at 320 K were used in this study as this is the closest to human body temperature. The ATLAS[4] database contains MD simulations for 1,516 PDB structures, each conducted using the CHARMM36m force field for 100 ns with three replicates. The Dynamic PDB[5] dataset includes over 10,000 molecular dynamics (MD) simulations of PDB structures, generated using the Amber-ff14SB force field. IDRome[6] contains conformational ensembles of human disordered regions generated via the coarse-grained residue-level CALVADOS model. All coarse-grained trajectories were downloaded and converted to all-atom trajectories using cg2all[7] (v1.5). While this conversion may result in structural artifacts and unrealistic conformations, we opted to use the converted all-atom trajectories as they provide more information, and deep learning models are generally robust to noise.

For IDRome, the first 10 frames were excluded as described in the original paper[6]; for mdCATH, no frame was discarded as the downloaded trajectories were recorded after the pre-equilibration phase; for the other trajectories, the first 20% of frames were discarded. Trajectories of the same protein were merged. All frames were aligned to the first frame based on Cα atoms using MDTraj[8]. mdCATH data was obtained in February 2025. Dynamic PDB data was obtained in March 2025, as this dataset was not fully open and under uploading, we obtained 756 proteins with 100 ns simulations. Other data were all obtained in January 2024.

#### Dynamic Properties Extraction from Structure Ensembles

GetContacts (https://getcontacts.github.io) was used to extract nine types of interactions from MD simulation trajectories: backbone-to-backbone hydrogen bonds, side-chain-to-backbone hydrogen bonds, side-chain-to-side-chain hydrogen bonds, salt bridges, Pi-cation, Pi-stacking, T-stacking, hydrophobic interactions, and van der Waals interactions. Default definitions of these interactions were used as described in https://getcontacts.github.io/interactions.html. For each residue pair, nine interaction frequencies were calculated, resulting in a matrix with the size of $L \times L \times 9$ for a protein of length $L$.

MDTraj[8] (v1.9.9) was used to extract residue-level properties. Surface area per residue was computed using mdtraj.shrake_rupley(mode='residue'), the mean and standard deviation of surface areas were calculated; Root mean square fluctuation (RMSF) was calculated using mdtraj.rmsf, as RMSF is related to simulation time, it was further normalized by dividing the max value in the protein; Eight-class secondary structures were determined using mdtraj.compute_dssp, the frequency of secondary structures was calculated; For dihedral angles, mdtraj.compute_phi, mdtraj.compute_psi, and mdtraj.compute_chi1 were employed to extract the *phi*, *psi*, and *chi1* angles, respectively. Dihedral angles were partitioned into 12 bins (30° intervals), and the percentage of frames falling into each bin was calculated for each residue. For residues don't have a specific angle (like terminal residues don't have *phi* or *psi*, Glycine don't have *chi1*), 1/12 was used. Collectively, this yielded a residue-level property matrix of size $L \times (2+1+8+3\times12)$ for a protein of length $L$, the dimension corresponds to surface area (2), normalized RMSF (1), secondary structure (8), and three dihedral angles distributions (3×12), respectively.

For the calculation of pairwise residue movement correlations, we first computed the covariance matrix for the *x*, *y*, and *z* coordinates of all Cα atoms:

$$C_{3L} = \frac{1}{p-1}\sum_{i=0}^{p}(X_i - \bar{X})\,(X_i - \bar{X})^T \quad (1)$$

Where $p$ is the number of frames in the ensemble, $X_i$ represents the positions (*x*, *y*, *z*) of all Cα atoms in frame $i$, and $\bar{X}$ is the mean position of the Cα atoms of all frames. The matrix $C_{3L}$ has a dimension of $3L \times 3L$ where $L$ is the number of Cα atoms (protein length).

To reduce this 3D covariance matrix to residue level, the trace over the spatial dimensions is taken:

$$C_L = Tr_{x,y,z}(C_{3L}) \tag{2}$$

Here, $C_L$ is the reduced covariance matrix, and the diagonal of $C_L$ corresponds to the squared fluctuation of *L* residues. The correlation matrix $R_L$ is then computed by normalizing the covariance matrix:

$$\sigma_i = \sqrt{C_{L,ii}} \tag{3}$$

$$R_{L,ij} = \frac{C_{L,ij}}{\sigma_i \sigma_j} \tag{4}$$

where $C_{L,ij}$ is the covariance between residues *i* and *j*, $\sigma_i$ and $\sigma_j$ are their respective standard deviations. This correlation matrix describes the linear relationship between the displacements of residue pairs, independent of their absolute motion magnitude.

#### Normal Mode Analysis

For normal mode analysis (NMA), PDB structures in ProteinFlow[9] were used. Structures containing gaps (missing residues in the middle) or exceeding 5,000 residues were excluded. Terminal missing residues were removed. For the 20230102_stable dataset, MMseqs2[10] cluster representatives at 90% sequence identity were used. For the 20231221_sabdab dataset, MMseqs2 cluster representatives at 100% identity were used. '*X*' was added between chains of complexes for MMseqs2 clustering.

NMA was conducted using the Gaussian Network Model (GNM)[11] and the Anisotropic Network Model (ANM)[12], both implemented in ProDy (v2.4.0)[13]. These models represent macromolecules as elastic node-and-spring networks, where Cα atoms are nodes, and springs connect residues within a defined cutoff distance. A distance-dependent spring force constant was applied as recommended in ProDy website (http://www.bahargroup.org/prody/tutorials/enm_analysis/gamma.html): for Cα atoms 10-15 Å apart, a unit force constant was used; for atoms 4-10 Å apart, a force constant twice as strong was used; and for atoms within 4 Å (i.e., connected residue pairs), a force constant 10 times stronger was employed. GNM, which models isotropic motion, was used to compute residue-level properties, while ANM, which captures anisotropic motion, was employed to calculate pairwise properties.

After building the elastic network (Kirchhoff matrix for GNM or Hessian matrix for ANM), normal modes were computed by eigenvalue decomposition. Eigenvalues ($\lambda_m$) and eigenvectors ($v_m$) describe the collective motions of residues in mode *m*. The individual contribution of each mode is the proportion of the inverse eigenvalue to all modes:

$$\frac{1/\lambda_m}{\sum_{k=1}^{M} 1/\lambda_k} \tag{5}$$

Where *M* is the number of total modes (*L-1* for GNM and *3L-6* for ANM, *L* is protein length). For model pre-training, modes of GNM and ANM were first ranked by contribution, then partitioned into three ranges separately. The ranges were selected such that the first set of modes accounts for ~33% of the dynamics, the second set for ~33–66%, and the final set for ~66–100%. This ensures that the slow, intermediate, and fast modes are separated. For evaluation in Figure 1F-G, certain number of low-frequency modes were used.

For each set of modes, the mean-square fluctuation (*MSF*) of each residue was calculated from GNM modes as:

$$MSF_i = \sum_m \frac{v_{mi}^2}{\lambda_m} \tag{6}$$

Where $v_{mi}$ is the eigenvector component corresponding to mode *m* and residue *i*, and $\lambda_m$ is the corresponding eigenvalue. This calculation was repeated for three mode ranges.

The residue covariance matrix was calculated using ANM modes. Firstly, the covariance matrix with the size of *3L×3L* was first computed as:

$$C_{3L} = \mathbf{V}\Lambda^{-1}\mathbf{V}^T \tag{7}$$

Where **V** is the matrix of eigenvectors and $\Lambda^{-1}$ is the inverse diagonal matrix of eigenvalues. To reduce the 3D covariance matrix to the residue-level, we used the same method as described in the calculation of pairwise residue movement correlations (equation 2-4).

#### Sequence Clustering

After addressing issues such as failed downloads, unreadable files, and processing errors (e.g., six trajectories from mdCATH had the error of atoms virtually overlapping reported by Mdtraj), we successfully extracted dynamic properties for 64,403 proteins (Table 1; excluding Dynamic PDB). To remove nearly duplicate proteins, we performed clustering using MMseqs2 with the parameters *--min-seq-id 1 -c 0.95 --cov-mode 0* (‘*X*’ was added between chains in complexes for clustering). We further selected sequences in each cluster with the following order: PED > GPCRmd > mdCATH > ATLAS > IDRome > ProteinFlow. This resulted in a final dataset of 61,959 proteins for model training and testing. We further clustered these sequences using MMseqs2 with the parameters *--min-seq-id 0.5, -c 0.8 --cov-mode 1*, yielding 43,193 clusters. We randomly selected 5% of the clusters as the test set, comprising 3,127 proteins, while the remaining 58,832 proteins formed the training set. The distribution of proteins across data sources is as follows: PED: 252 (train), 17 (test); GPCRmd: 107 (train), 5 (test); mdCATH: 5,067 (train), 312 (test); ATLAS: 1,361 (train), 71 (test); IDRome: 26,502 (train), 1,360 (test); ProteinFlow: 25,543 (train), 1,362 (test).

Dynamic PDB was used as an independent test set because its trajectories were generated using a different force field (Amber) compared to the other datasets (CHARMM and coarse-grained). Proteins in Dynamic PDB dataset were searched against SeqDance/ESMDance training set using MMseqs2 search with the parameters: *-s 7 -a 1*, only 29 proteins without homologs (over 50% identity and 80% coverage) in the training set were analyzed in Figure 2.

### Figures

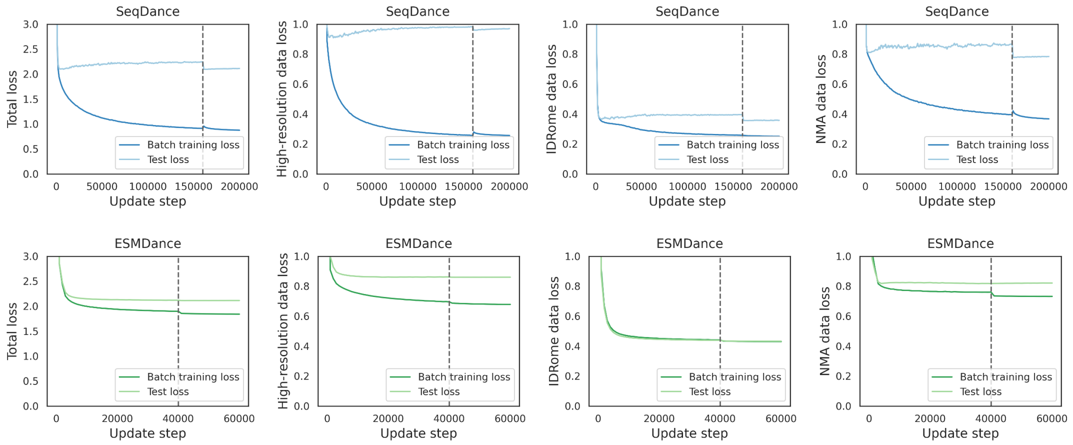


**Figure S1. Training loss and test loss curves for SeqDance and ESMDance.**

Models were trained with a maximum sequence length of 256 before the dashed line and 1,024 after the dashed line. Test losses were calculated using the first 1,024 residues for long proteins. The training and test sets were split at 50% sequence identity. Loss values from each data source were normalized such that the baseline model loss was set to 1, resulting in the total loss of 3 (the baseline model predicts the mean for each property of each protein, see Methods for details).

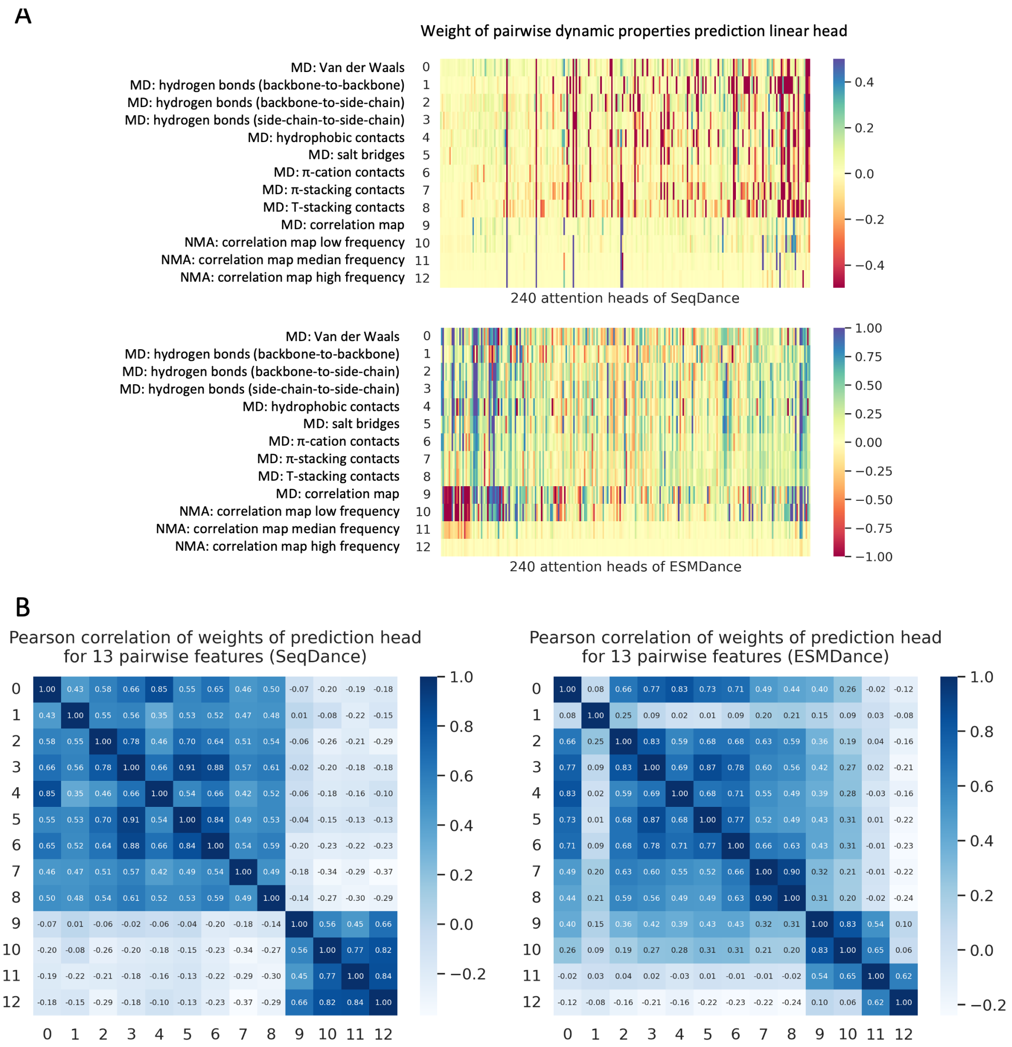


**Figure S2. Analysis of weights of the pairwise property prediction linear layer.**

**A**. Heatmaps of weights of linear layers that use 240 attention maps to predict pairwise properties. For a protein of length *L*, the pairwise properties' dimension is $L \times L \times 13$. The input for the pairwise property prediction head consists of pairwise embeddings derived from residue embeddings and attention maps (dimension $L \times L \times 240$). Only weight for attention maps shown here. **B**. Pearson correlation between weights of prediction heads for different properties.

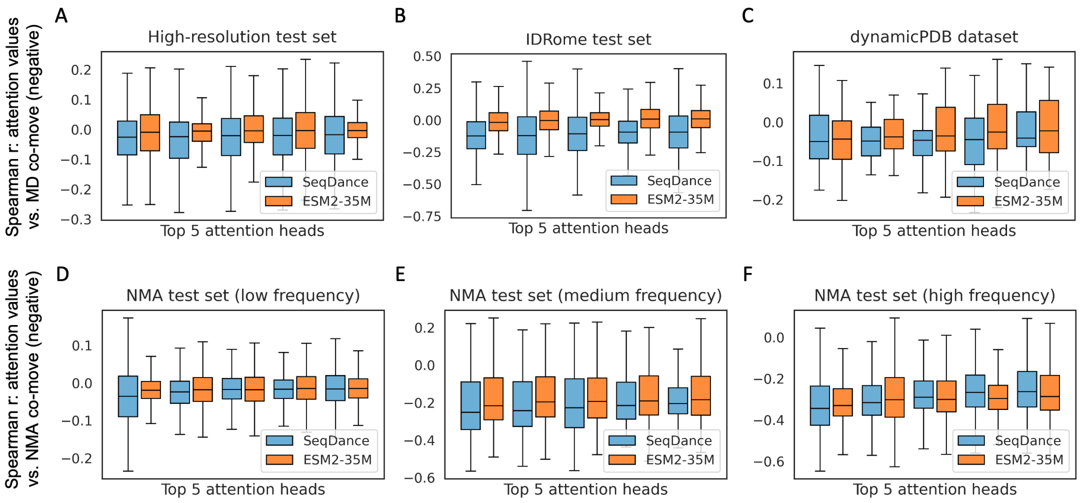


**Figure S3. Analysis of attention and negatively correlated residue pairs.**

For each attention head, Spearman correlation was calculated between attention values and pairwise dynamic properties, top five heads of SeqDance and ESM2-35M are shown. Only negatively correlated residue pairs are included, negative correlations indicate better capture of these properties. Boxplots show the distribution of Spearman correlations of test data, The box extends from the first quartile to the third quartile of the data, with a line at the median. The whiskers extend from the box to the farthest data point lying within 1.5 times the inter-quartile range from the box.

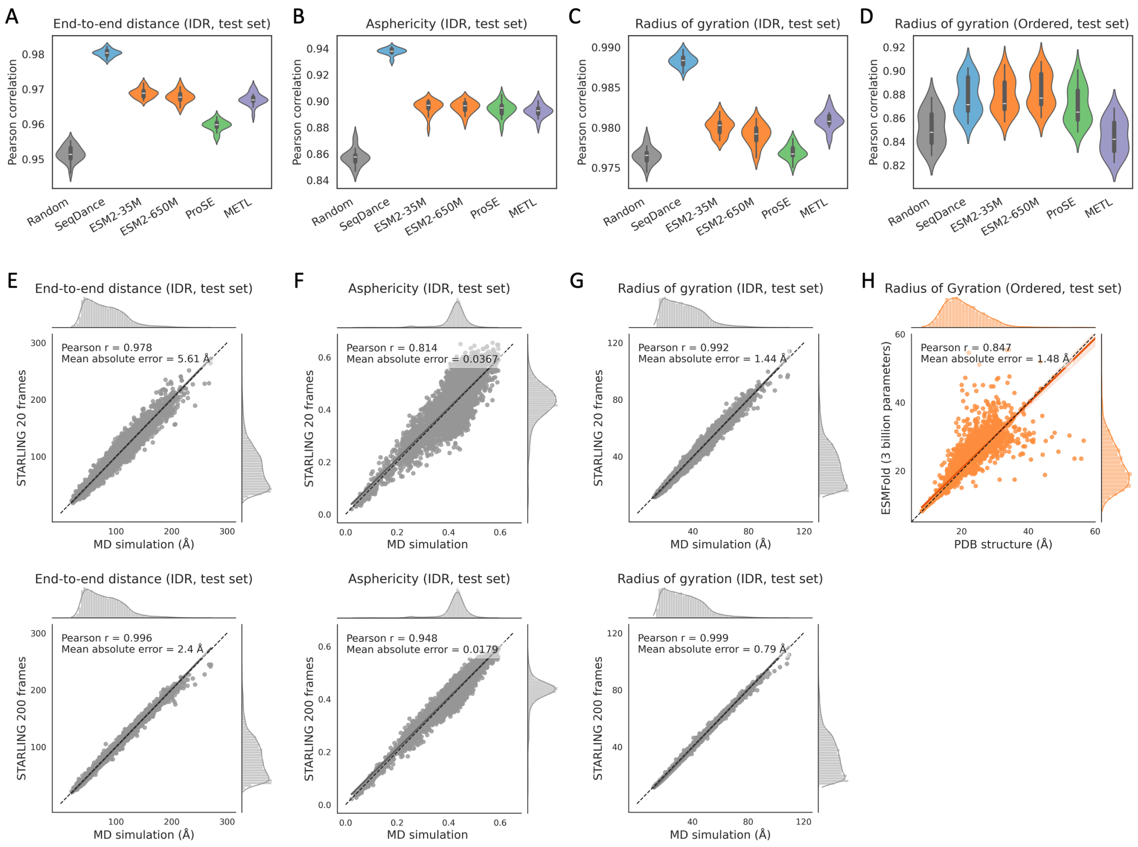


**Figure S4. SeqDance's embeddings encode global conformational properties.**

**A-D**. Performance (Pearson correlation) comparison of embeddings from SeqDance, METL, ProSE, and ESM2 in predicting the end-to-end distance of disordered regions (**A**), asphericity of disordered regions (**B**), radius of gyration of disordered regions (**C**) and ordered proteins (**D**). The training and test split was 6:4 with a 20% sequence identity cutoff. Linear regression model was trained to predict conformational property using the first 200 principal components of embeddings from each method as input. The results presented are the distributions of 20 independent repeats. **E-G**. Comparison of conformational properties of test IDRs obtained from coarse-grained MD simulations versus STARLING-generated ensembles. The upper panels show results using 20 STARLING-generated conformations, and the lower panels show results using 200 conformations. **H**. Comparison of the radius of gyration of test ordered proteins derived from PDB structures versus ESMFold predictions. In panels **E–H**, each dot represents a test protein; the grey/orange line indicates the linear fit, and the dashed black line represents the line of equality.

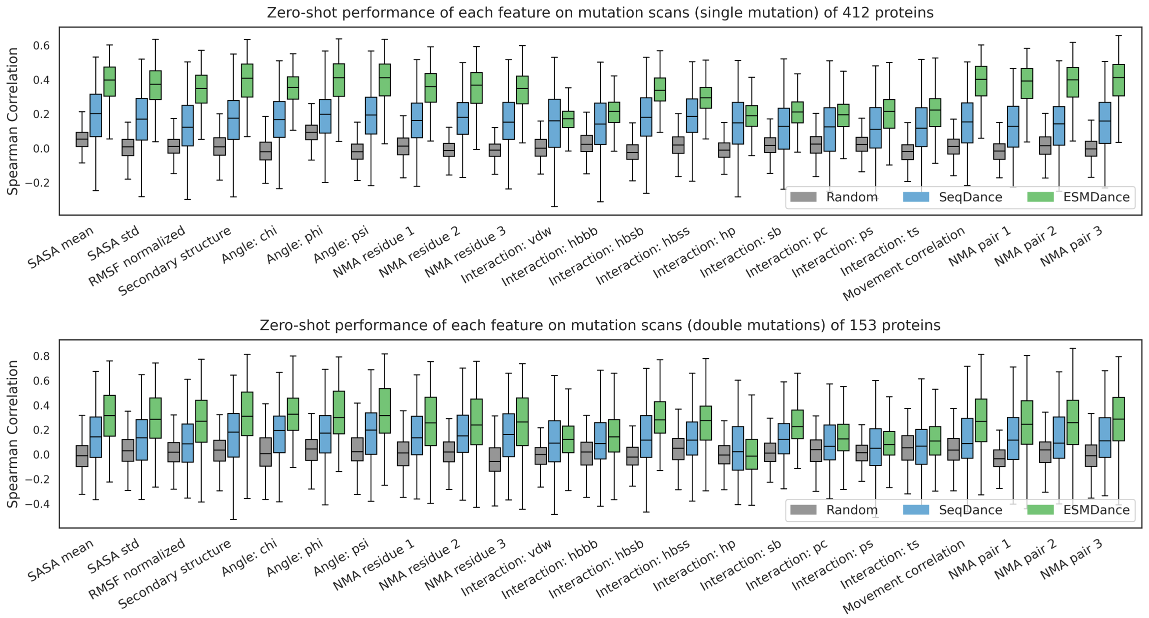


**Figure S5. Zero-shot prediction performance for mutation effect on stability using individual dynamic properties.**

Distribution of zero-shot performance (Spearman correlation) of different dynamic properties from the random model, SeqDance, and ESMDance. The random model (color gray) represents the randomly initialized model prior to SeqDance pre-training. Single mutations and double mutations are evaluated separately. While all 412 proteins have single mutation scanning experiments, only 153 of them have double mutation scanning experiments.

SASA mean, std: mean and standard deviation of solvent-accessible surface area. NMA properties 1, 2, 3: properties calculated from low-, median-, and high-frequency normal modes. vdw: van der Waals interaction; hbbb: backbone-to-backbone hydrogen bond; hbsb: side-chain-to-backbone hydrogen bond; hbss: side-chain-to-side-chain hydrogen bond; hp: hydrophobic interaction; sb: salt bridge; pc: Pi-cation interaction; ps: Pi-stacking interaction; ts: T-stacking interaction.

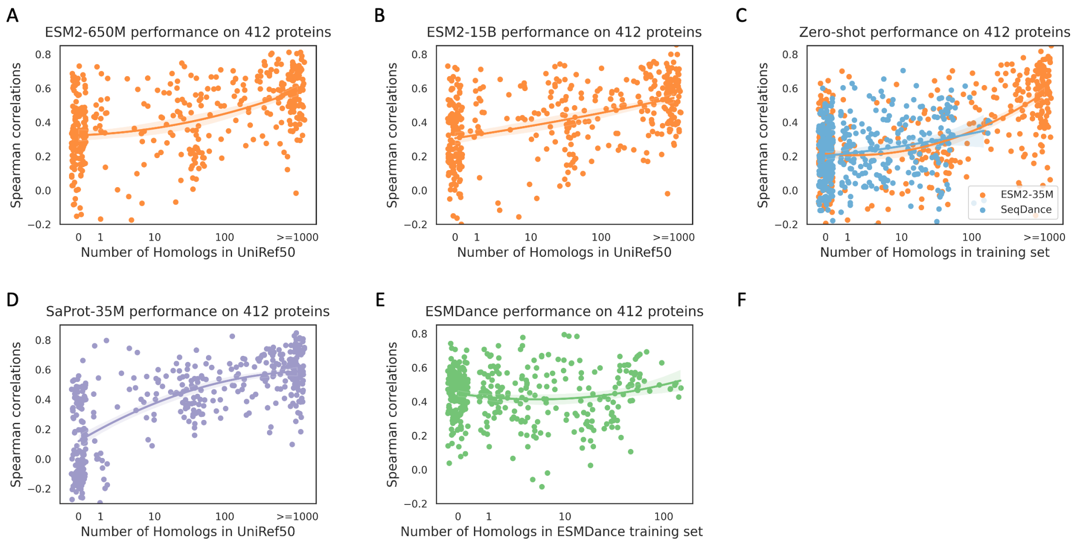


**Figure S6. Zero-shot prediction of mutation effect on stability.**

Relationship between zero-shot performance and the number of homologs (defined as over 20% sequence identity and 50% coverage) in UniRef50 for ESM2-650M (**A**), -15B (**B**), and SaProt-35M (**D**). **C**. Relationship between zero-shot performance and the number of homologs in the SeqDance training set for SeqDance and in UniRef50 for ESM2-35M. This plot is the combination of main Figure 4D and 4E. **E**. Relationship between ESMDance zero-shot performance and the number of homologs in the ESMDance training set.

The line represents a polynomial regression of order two, with the shaded area indicating the 95% confidence interval, the x-axis shows the log-scaled number of homologs, with small random noises added to the x values to reduce overlap.

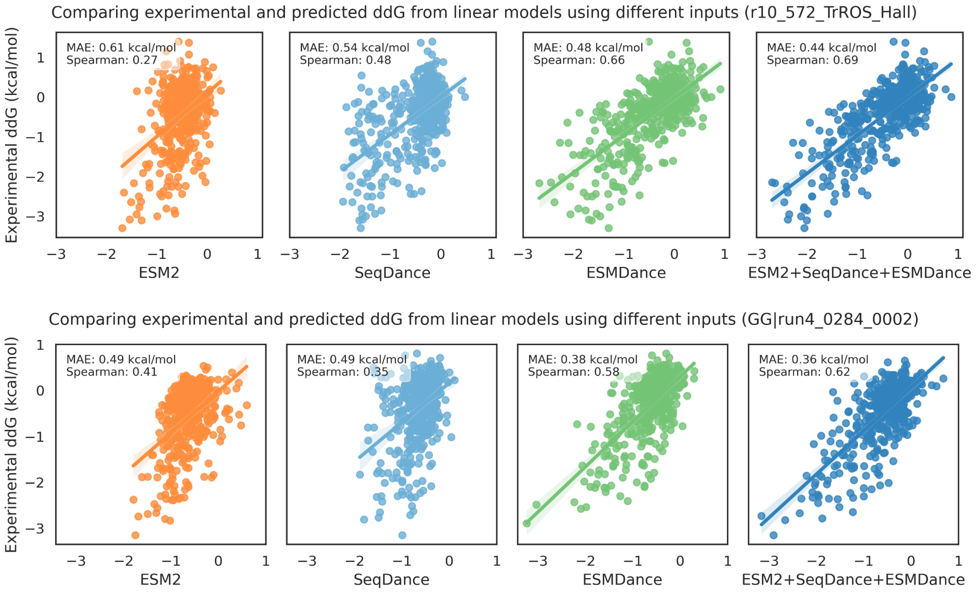


**Figure S7. Linear regression models predicted mutation effect of two designed proteins.**

Linear regression models were trained to align zero-shot predictions from these models with experimental ddG. For each protein, 50% of mutations were randomly sampled as training set, and the remaining 50% were used as test, the predicted and experimental ddG values on test mutations were shown here. Zero-shot predictions from three ESM2 models (35M, 650M, 15B) and/or 23 SeqDance/ESMDance-predicted dynamic property changes were used as input. MAE: mean absolute error.

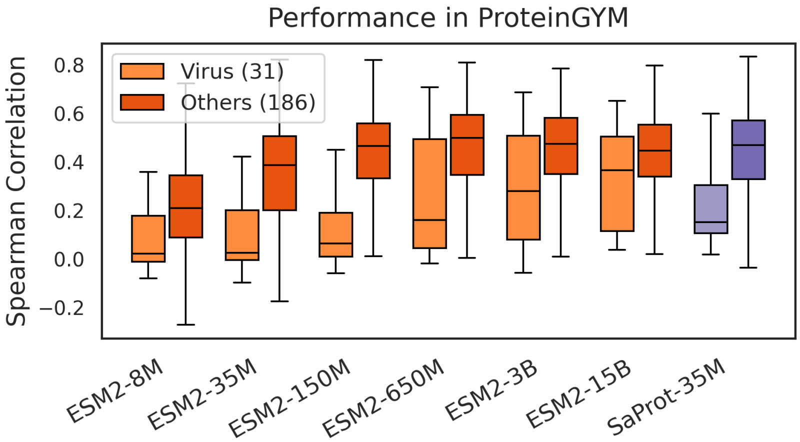


**Figure S8. Zero-shot prediction of mutation effect for viral proteins and others.**

Zero-shot performance of ESM2 models and SaProt-35M for viral proteins and other proteins. The Spearman correlations for models were downloaded from ProteinGYM.

### Tables

#### Table S1: residue-level and pairwise dynamic properties

| Method | Feature | Dimension | Description | Package |
|---|---|---|---|---|
| NMA | Correlation map | *L×L×3* | Normal modes accounting for 33%, 66%, and 100% of overall dynamics in ANM (for correlation map) or GNM (for residue fluctuation) | ProDy |
| | Residue fluctuation | *L×3* | | |
| MD | Correlation map | *L×L×1* | Correlation of Cα movement | mdtraj |
| | Interaction map | *L×L×9* | Hydrogen bonds (side-chain-to-side-chain, backbone-to-backbone, backbone-to-side-chain), salt bridges, hydrophobic contacts, π-cation contacts, π-stacking contacts, T-stacking contacts, Van der Waals | GetContacts |
| | Residue fluctuation | *L×1* | Normalized root mean squared fluctuation (RMSF) | mdtraj |
| | Surface Area | *L×2* | Mean and standard deviation | mdtraj |
| | Secondary Structure | *L×8* | Percentage of eight DSSP assignments | mdtraj |
| | Dihedral angles: *phi*, *psi*, *chi1* | *L×(3×12)* | Percentages in 12 angle ranges | mdtraj |

NMA: normal mode analysis

MD: molecular dynamics

ANM: Anisotropic Network Model

GNM: Gaussian Network Model

*L*: protein length

**Table S2: performances of predicting conformational properties using full embeddings.**

| | | AAindex | Random | SeqDance | ESM2-35M | ESM2-650M | ProSE | METL |
|---|---|---|---|---|---|---|---|---|
| Dim of embeddings | | 563 | 480 | 480 | 480 | 1280 | **6165** | 512 |
| End-to-end distance (IDR) | MAE (Å) ↓ | 9.016 | 5.198 | **3.332** | 4.594 | 3.791 | 3.813 | 4.886 |
| | Pearson r ↑ | 0.873 | 0.959 | **0.986** | 0.977 | 0.983 | 0.985 | 0.975 |
| Asphericity (IDR) | MAE ↓ | 0.027 | 0.02 | **0.013** | 0.018 | 0.015 | 0.015 | 0.019 |
| | Pearson r ↑ | 0.702 | 0.877 | **0.949** | 0.917 | 0.934 | 0.945 | 0.912 |
| Radius of gyration (IDR) | MAE (Å) ↓ | 2.879 | 1.525 | **1.027** | 1.441 | 1.19 | 1.11 | 1.48 |
| | Pearson r ↑ | 0.928 | 0.981 | 0.991 | 0.985 | 0.989 | **0.992** | 0.986 |
| Radius of gyration (ordered) | MAE (Å) ↓ | 1.929 | 1.954 | **1.651** | 1.701 | 1.882 | 4.353 | 2.001 |
| | Pearson r ↑ | 0.844 | 0.845 | **0.875** | 0.874 | 0.868 | 0.625 | 0.84 |

The values in the table are the mean performance on test sets of 20 independent experiments.

MAE: mean absolute error.

IDR: intrinsically disordered region.

AAindex (https://www.genome.jp/aaindex/): the mean of 563 chemical and physical properties of all residues in the protein.

Random: the randomly initialized model prior to SeqDance pre-training.

ProSE embeddings are the concatenation of embeddings from all layers, as suggested in its GitHub code. It seems that ProSE overfits on the training set of ordered proteins.

**Table S3: zero-shot performance (Spearman correlation) of different combinations of relative changes of dynamic properties on 412 proteins**

| Relative changes | Combination methods | SeqDance mean | SeqDance median | ESMDance mean | ESMDance median |
|---|---|---|---|---|---|
| Raw | Mean | 0.2146 | 0.2157 | 0.4321 | 0.4491 |
| | Max | 0.1772 | 0.1861 | 0.3763 | 0.3884 |
| | Weighted mean | 0.1751 | 0.1756 | 0.3955 | 0.4157 |
| | Geometric mean | 0.2160 | 0.2210 | 0.4141 | 0.4303 |
| Quantile normalization | Mean | 0.2293 | **0.2407** | 0.4313 | 0.4497 |
| | Max | 0.2031 | 0.2012 | 0.2794 | 0.2857 |
| | Weighted mean | 0.2237 | 0.2272 | 0.3939 | 0.4078 |
| | Geometric mean | **0.2317** | 0.2383 | **0.4383** | **0.4580** |

### SI References


1. Ghafouri, H. et al. PED in 2024: improving the community deposition of structural ensembles for intrinsically disordered proteins. Nucleic Acids Res 52, D536–D544 (2024).

2. Rodriguez-Espigares, I. et al. GPCRmd uncovers the dynamics of the 3D-GPCRome. Nat Methods 17, 777–787 (2020).

3. Mirarchi, A., Giorgino, T. & De Fabritiis, G. mdCATH: A Large-Scale MD Dataset for Data-Driven Computational Biophysics. Sci Data 11, 1299 (2024).

4. Vander Meersche, Y., Cretin, G., Gheeraert, A., Gelly, J. C. & Galochkina, T. ATLAS: protein flexibility description from atomistic molecular dynamics simulations. Nucleic Acids Res 52, D384–D392 (2024).

5. Liu, C. et al. Dynamic PDB: A New Dataset and a SE(3) Model Extension by Integrating Dynamic Behaviors and Physical Properties in Protein Structures. Preprint at https://doi.org/10.48550/arXiv.2408.12413 (2024).

6. Tesei, G. et al. Conformational ensembles of the human intrinsically disordered proteome. Nature https://doi.org/10.1038/s41586-023-07004-5 (2024) doi:10.1038/s41586-023-07004-5.

7. Pang, Y. T., Yang, L. & Gumbart, J. C. From simple to complex: Reconstructing all-atom structures from coarse-grained models using cg2all. Structure 32, 5–7 (2024).

8. McGibbon, R. T. et al. MDTraj: A Modern Open Library for the Analysis of Molecular Dynamics Trajectories. Biophys J 109, 1528–32 (2015).

9. Mi, Y., Marcu, S.-B., Yallapragada, V. V. B. & Tabirca, S. ProteinFlow: An advanced framework for feature engineering in protein data analysis. Biotechnol. Bioeng. 121, 3563–3571 (2024).

10. Steinegger, M. & Soding, J. MMseqs2 enables sensitive protein sequence searching for the analysis of massive data sets. Nat Biotechnol 35, 1026–1028 (2017).

11. Haliloglu, T., Bahar, I. & Erman, B. Gaussian Dynamics of Folded Proteins. Phys. Rev. Lett. 79, 3090–3093 (1997).

12. Atilgan, A. R. et al. Anisotropy of fluctuation dynamics of proteins with an elastic network model. Biophys J 80, 505–15 (2001).

13. Bakan, A., Meireles, L. M. & Bahar, I. ProDy: Protein Dynamics Inferred from Theory and Experiments. Bioinformatics 27, 1575–1577 (2011).
